# Ten years of unpredictable chronic stress in zebrafish: a systematic review and meta-analysis

**DOI:** 10.1101/2022.12.12.520151

**Authors:** Matheus Gallas-Lopes, Leonardo M. Bastos, Radharani Benvenutti, Alana C. Panzenhagen, Angelo Piato, Ana P. Herrmann

**Affiliations:** Brazilian Reproducibility Initiative in preclinical Systematic Review and Meta-Analysis (BRISA) Collaboration, Rio de Janeiro, Brazil; Laboratório de Neurobiologia e Psicofarmacologia Experimental (PsychoLab), Departamento de Farmacologia, Instituto de Ciências Básicas da Saúde, Universidade Federal do Rio Grande do Sul, Rua Ramiro Barcelos 2600/430, Porto Alegre, Rio Grande do Sul, 90035-003, Brazil; Laboratório de Psicofarmacologia e Comportamento (LAPCOM), Departamento de Farmacologia, Instituto de Ciências Básicas da Saúde, Universidade Federal do Rio Grande do Sul (UFRGS), Rua Ramiro Barcelos 2600/411, Porto Alegre, RS, 90035- 003, Brazil; Department of Physiology and Medical Physics, Royal College of Surgeons in Ireland, 123 St. Stephen’s Green, Dublin, Ireland; Programa de Pós-Graduação em Ciências Biológicas: Bioquímica, Instituto de Ciências Básicas da Saúde, Universidade Federal do Rio Grande do Sul (UFRGS), Rua Ramiro Barcelos 2600, Porto Alegre, RS, 90035-003, Brazil

**Keywords:** Unpredictable chronic stress, *Danio rerio*, animal model, anxiety, locomotor function, social behaviour, cortisol, systematic review, meta-analysis, depression

## Abstract

The zebrafish (*Danio rerio*) is a model animal that is being increasingly used in neuroscience research. A decade ago, the first study on unpredictable chronic stress (UCS) in zebrafish was published, inspired by protocols established for rodents in the early 1980’s. Since then, several studies have been published by different groups, in some cases with conflicting results. We conducted a systematic review to identify studies evaluating the effects of UCS in zebrafish and meta-analytically synthetized the data of neurobehavioral outcomes and relevant biomarkers. Literature searches were performed in three databases (PubMed, Scopus, and Web of Science) and a two-step screening process based on inclusion/exclusion criteria. The included studies underwent extraction of qualitative and quantitative data, as well as risk of bias assessment. Outcomes of included studies (*n =* 38) were grouped into anxiety/fear-related behaviour, locomotor function, social behaviour, or cortisol level domains. UCS increased anxiety/fear-related behaviour and cortisol levels while decreased locomotor function, but a significant summary effect was not observed for social behaviour. Despite including a significant number of studies, the high heterogeneity and the methodological and reporting problems evidenced in the risk of bias analysis make it difficult to assess the internal validity of most studies and the overall validity of the model. Our review thus evidences the need to conduct well-designed experiments to better evaluate the effects of UCS on the behaviour of zebrafish.

## I. INTRODUCTION

The origins of the unpredictable chronic stress (UCS) protocol go back to the early 1980s, when researchers proposed the chronic administration of a variety of stressors to rodents as a way to induce behavioural alterations relevant to the study of depression (Katz & Hersh, 1981; Katz, Roth & Carroll, 1981; Katz, 1982; Willner *et al*., 1987). Construct, face, and predictive validities of this model are supported by many studies showing that rodents exposed to the UCS protocol develop anhedonia-like behaviour, cognitive deficits, hormonal and neurochemical imbalances, weight loss, and other changes that can be reversed by using antidepressant treatments (Willner, 1997). Given its translational potential, there has been an exponential growth in the implementation of this protocol across laboratories as it has become an important tool for the study of the neurobiological basis of depression and antidepressant action (Willner, 2017a; Nollet, 2021).

Whereas this intervention became popular, researchers started adapting the UCS protocol and reports of controversial data and reproducibility problems have also increased (Strekalova & Steinbusch, 2009; Willner, 2017b; Antoniuk *et al*., 2019). The protocol has been largely criticized for its lack of reliability as many known elements such as the training level of experimenters, the duration of the protocol, and animal characteristics (species, strain, sex, and others) can introduce variability and influence the results (Willner, 2017b). Apart from that, even with heterogeneous protocols, UCS- induced behavioural and physiological alterations have been replicated within and between labs, adding to the internal and external validity of the model.

More than a decade ago, researchers made an effort to transpose this intervention for studies using zebrafish (*Danio rerio* Hamilton, 1822), an emerging model animal in the field of neuroscience at the time (Piato *et al*., 2011). Cross-species approaches are important tools to evaluate the validity of an intervention, and translating the UCS protocol to zebrafish can help reduce species-specific biases originating from studies conducted solely with rodents (Maximino *et al*., 2015; Weber- Stadlbauer & Meyer, 2019). In zebrafish, this protocol is also able to induce anxiety- like behaviour and alterations in outcomes such as locomotion, cognition, sociability, cortisol levels, and in some defence mechanisms against oxidative damage (Piato *et al*., 2011; Marcon *et al*., 2016, 2018; Bertelli *et al*., 2021). But just as in the experiments carried out with rats and mice, the heterogeneity between protocols established in each laboratory has grown throughout the years as investigators needed to adapt the procedures to different facilities or to the outcomes of interest. This culminated in the publication of many discrepant results for key outcomes to understand the impacts of UCS, like social behaviour, which was shown to be altered in opposing directions depending on the duration of the protocol (Piato *et al*., 2011), or not altered at all (Golla, Østby & Kermen, 2020; Bertelli *et al*., 2021).

Aiming to estimate the overall validity and to summarise the evidence regarding the effects of UCS on behavioural and biochemical outcomes relevant to the study of psychiatric disorders, we conducted a systematic review and meta-analysis of the available scientific literature using zebrafish. We analysed the evolution of this intervention in the first ten years of its use, qualitatively describing the published studies, establishing the direction and magnitude of the effect of chronic stress on neurobehavioural and neurochemical parameters, detecting effect moderators, and evaluating the impact of bias arising from methodological conduct, reporting quality, and selective publication.

## II. METHODS

A protocol for conducting this review was registered on Open Science Framework prior to the screening of records and data collection. Preregistration is available at https://osf.io/9rvyn (Gallas-Lopes *et al*., 2021). The reporting of this study complies with the Preferred Reporting Items for Systematic Reviews and Meta-Analyses (PRISMA) guidelines (Page *et al*., 2021).

### 1 Search strategy

Searches were conducted in three bibliographic databases: PubMed, Scopus, and Web of Science. The search strategy was designed to include broad terms that describe the intervention (UCS protocol) and the desired population (zebrafish). The complete query for each database can be found at https://osf.io/9rvyn (Gallas-Lopes *et al*., 2021). There were no language or date restrictions. The first search was performed on the 10^th^ of July, 2021, with an update search carried out on the 26^th^ of October, 2021. The bibliographic data acquired were imported to Rayyan software (Ouzzani *et al*., 2016), where duplicates were detected and removed by one of the investigators (MGL). The reference lists of the included studies were also screened in order to detect additional relevant articles.

### 2 Eligibility screening

After the removal of duplicates, the selection of eligible studies was conducted using Rayyan software in a two-step process based, initially, on title and abstract, followed by a full-text analysis. The screening of each record was performed by two independent investigators (MGL and LMB or RB) and disagreements were resolved by a third investigator (APH). Peer-reviewed articles were eligible for inclusion if they had an appropriate control group and assessed the effects of unpredictable chronic stress in zebrafish (any strain or developmental stage) on any of the following domains of interest: morphometric measures, locomotor function, sensory function, learning and memory, social behaviour, reproductive behaviour, anxiety/fear-related behaviour, circadian cycle-related behaviour, and neurochemical or peripheral biomarkers (e.g., cortisol, cytokines, and oxidative stress).

In the first screening stage (title and abstract), studies were excluded based on the following reasons: (1) design: not an original primary study (e.g., review, commentary, conference proceedings, and corrections); (2) population: studies using other species than zebrafish (*Danio rerio*) or studies that did not use any animal; (3) intervention: non-interventional studies or studies using other interventions than unpredictable chronicle stress (e.g., acute stress (stressed only once) and repetitive or predictable stress (chronic stress using only a single stressor multiple times)). In the second stage (full-text screening), the remaining articles were assessed for exclusion based on the same reasons considered in the first stage plus the following additional reasons: (4) comparison: studies without an adequate control group; (5) outcome: studies that did not evaluate any of the target outcomes. All Rayyan files with investigators’ decisions are available at the study repository in Open Science Framework (https://osf.io/j2zva/), section “Eligibility screening archives”.

### 3 Data extraction

Data extraction from included studies was conducted by two independent investigators (MGL and LMB or RB) and disagreements were resolved by a third investigator (APH). Whenever available, the exact information and values were extracted directly from text or tables. Otherwise, WebPlotDigitizer software (v4.5, Rohatgi, A., Pacifica, CA, USA, https://automeris.io/WebPlotDigitizer) was used to manually estimate numbers from the graphs. In cases of lacking or dubious information, investigators attempted to contact via e-mail the corresponding author of the study in two separate attempts, at least two weeks apart.

The following characteristics were extracted: (1) study characteristics: study title, digital object identifier (DOI), first and last authors, last author’s institutional affiliation, and year of publication; (2) animal model characteristics: strain, sex, animal source (supplier of the animals used to develop the experiments), the total number of animals used, and the developmental stages during stress induction and outcome assessment; (3) UCS protocol characteristics: the number of different stressors, stress sessions per day, stress sessions in total, the duration of the stress protocol in days, and the time in days between the end of UCS protocol and outcome assessment; (4) test characteristics: experiment identification (to annotate whether the tests conducted within the same study used different sets of animals), the type of the test, test duration, habituation phase (whether the animals were subjected to an habituation phase in the experimental apparatus prior to the test), the category of the measured variable, and the measured variable. Co-authorship networks were constructed using VOSviewer software version 1.6.18 (https://www.vosviewer.com) (van Eck & Waltman, 2007,

2010).

Outcome data were extracted for each of the variables within the domains of interest. The measure of central tendency and the number of animals (n) were extracted for the control and UCS groups along with the standard deviation (SD) or standard error (SEM) when the mean value was expressed, or the interquartile range (IQR) when data were expressed as the median value. Whenever sample size was reported as a range instead of the exact number of animals in each group, the lowest value was extracted. If the study reported the SEM, SD was calculated by multiplying SEM by the square root of the sample size (SD = SEM ∗ √n).

### 4 Risk of bias and reporting quality

In order to evaluate the quality of included studies, the risk of bias assessment was conducted by two independent investigators (MGL and LMB or RB) for each paper, and disagreements were resolved by a third investigator (APH). The analysis was conducted based on the SYRCLE’s risk of bias tool for animal studies (Hooijmans *et al*., 2014) with adaptations to better suit the model animal and the intervention of interest. The following items were evaluated for methodological quality: (1) description of random allocation of animals; (2) description of baseline characteristics; (3) description of random housing conditions during the experiments; (4) description of random selection for outcome assessment; (5) description of blinding methods for outcome assessment; (6) incomplete outcome data; (7) selective outcome reporting. Additionally, four other items were evaluated by the investigators to assess the overall reporting quality of the studies based on a set of reporting standards for rigorous study design (Landis *et al*., 2012): (8.1) mention of any randomization process; (8.2) sample size estimation; (8.3) mention of inclusion/exclusion criteria; (8.4) mention of any process to ensure blinding during the experiments. For methodological quality, each item was scored with a “Yes” for low risk of bias, “No” for a high risk of bias or “Unclear” when it was not possible to estimate the risk of bias based on the information provided. Items regarding reporting quality were scored with only “Yes’’ or “No”, meaning high or low risk of bias, respectively. A complete guide for assessing the risk of bias associated with each of the items in this review is available at https://osf.io/sdpwb.

Risk of bias plots were created using *robvis* (McGuinness & Higgins, 2021).

### 5 Meta-analysis

Studies were grouped based on the domains of interest (anxiety/fear-related behaviour, locomotor function, social behaviour, or cortisol levels), and a meta-analysis was performed for each group. When a study reported multiple outcomes for the same domain, only one outcome of interest was chosen for the meta-analysis based on a rank of frequency developed by one of the investigators (MGL). Tests and variables within each test were ranked prior to data extraction, and the most frequent in the rank was included in the meta-analysis. The ranking is available at https://osf.io/rvn8b. A minimum of five studies were required for each domain in order to conduct a meta-analysis, as established a priori in our protocol (Gallas-Lopes *et al*., 2021).

The sample size of the control group was divided by the number of comparisons and rounded down whenever two or more experimental groups shared the same control (Vesterinen *et al*., 2014). When outcomes were analysed across time, the last point was selected for analysis. When animals were subjected to experiments at different time points following the end of the UCS protocol, the outcomes assessed closest to the end of the protocol were chosen. Effect sizes were “flipped” (multiplied by minus one) when needed to adjust the direction of the effect for specific behavioural traits in order to properly interpret the effects of UCS. Studies that only reported outcomes as the median value and interquartile range were excluded from the analyses along with studies with incomplete data (e.g., lacking sample sizes, SD, and SEM) when contact with the authors was unsuccessful.

Effects sizes were determined with standardised mean differences (SMD) using Hedge’s G method. Analyses were conducted using R Project for Statistical Computing with packages *meta* (Balduzzi, Rücker & Schwarzer, 2019) (https://cran.r-project.org/package=meta) and *ggplot2* (Wilkinson, 2011) following Hedge’s random effects model given the anticipated heterogeneity between studies. Values for SMD were reported with 95% confidence intervals. Heterogeneity between studies was estimated using I² (Higgins & Thompson, 2002), τ^2^, and Cochran’s Q (Cochran, 1954) tests. Heterogeneity variance (τ^2^) was estimated using the restricted maximum likelihood estimator (Viechtbauer, 2005; Veroniki *et al*., 2016). The confidence intervals around pooled effects were corrected using Knapp-Hartung adjustments (Knapp & Hartung, 2003). Values of 25%, 50%, and 75% were considered as representing low, moderate, and high heterogeneity, respectively for I², and a *p*-value ≤ 0.1 was considered significant for Cochran’s Q. Prediction intervals were estimated to present the range of effects expected for future studies (Higgins *et al*., 2019). Furthermore, a subgroup meta-analysis was performed to evaluate if the duration of the UCS protocol was a potential source of heterogeneity. Studies were grouped into two categories: those with up to 7 days of UCS protocol and those with more than 7 days. Subgroup analysis was only performed when there were at least five unique studies for each subgroup. A *p* ≤ 0.1 was considered significant for subgroup differences (Richardson, Garner & Donegan, 2019).

Publication bias was investigated by generating funnel plots and performing Duval and Tweedie’s trim and fill analysis (Duval & Tweedie, 2000) and Egger’s regression test (Egger *et al*., 1997). Analyses were only conducted when at least five studies were available within a given domain for funnel plots and at least ten studies for the regression test. A *p-*value < 0.1 was considered significant for the regression test.

### 6 Sensitivity analysis

A sensitivity analysis was conducted to assess if any experimental or methodological difference between studies was distorting the main effect found in the meta-analysis. Analyses were conducted following the jackknife method (Miller, 1974) and by excluding studies presenting a significant risk of bias, defined as either a high risk of bias in one of the main items evaluating methodological quality (items 1 to 7), or an unclear risk of bias in five or more of the same items. A minimum of three comparisons were required for each domain in order to conduct a sensitivity analysis.

## III. RESULTS

### 1 Search results

From the search in the selected databases, 420 records were retrieved altogether. Following the removal of duplicates, 206 records were screened for eligibility based on title and abstract. After the first screening phase, 58 studies remained to be assessed based on full text, and 38 met the criteria and were included in the review (Fig. 1). Out of the studies included in the review, 34 were collected from the first database search on the 10^th^ of July 2021, and four additional studies were identified in the second search on the 26^th^ of October 2021. No extra studies were identified by reference list screening. Most of the records sought for inclusion in either stage of screening were excluded because they did not meet the criteria set for the intervention (*n* = 89), followed by the population of interest (*n* = 42), and the design of the study (*n* = 37). Three studies were excluded from the quantitative analyses because the minimum number of studies to perform a meta-analysis was not reached for the outcomes reported (Zimmermann *et al*., 2016; Jayamurali & Govindarajulu, 2017; Marcon *et al*., 2018), and four studies were excluded because of missing information (Huang, Butler & Lubin, 2019; Zhang *et al*., 2021; Kirsten *et al*., 2021; Demin *et al*., 2021). This resulted in 31 studies included in the quantitative synthesis.

**Fig. 1.**
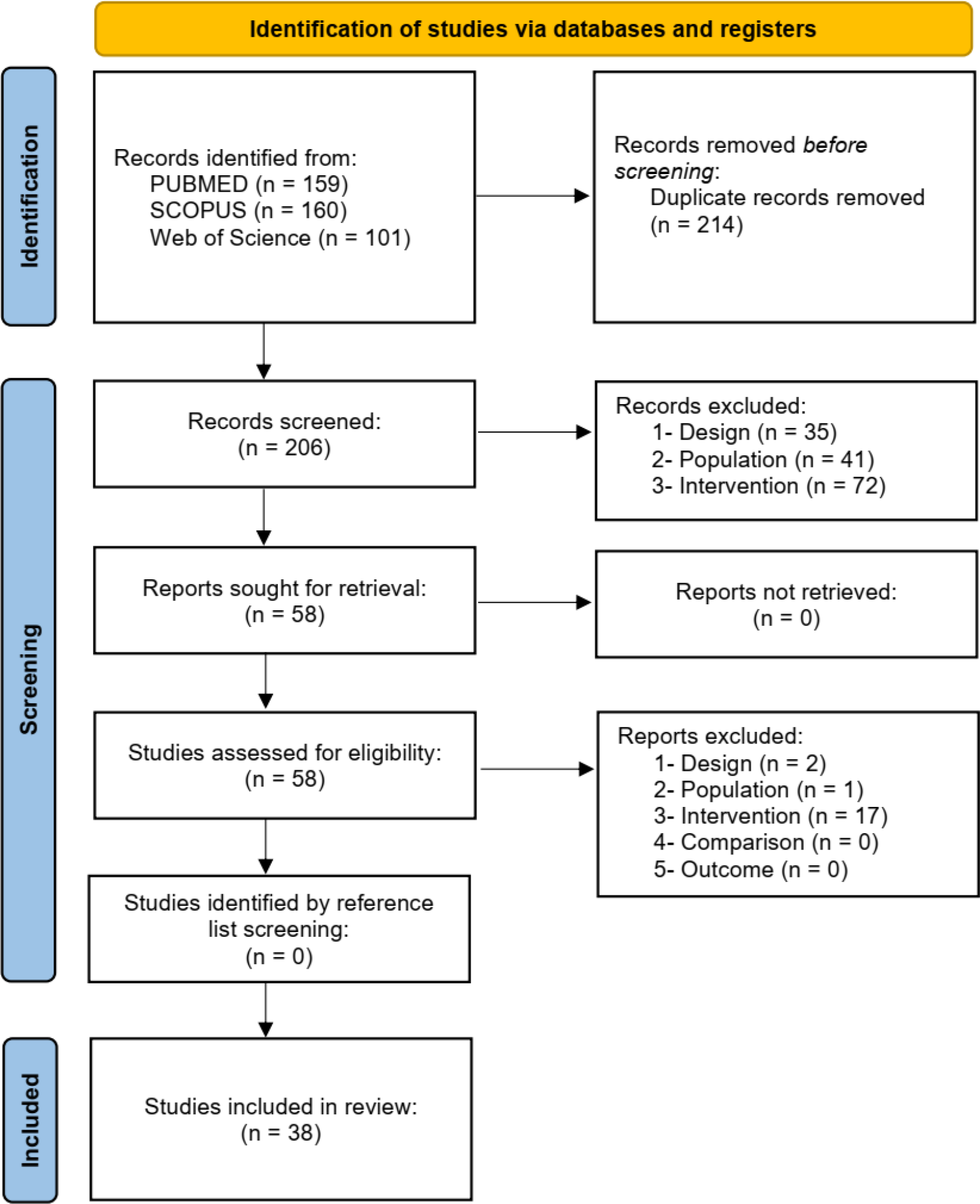
Flowchart diagram of the collection of studies and selection process.

### 2 Study characteristics

As expected, the protocols implemented by each research group varied significantly. The duration of the stress protocol ranged between 3 and 77 days, with 15 studies (39.5%) implementing UCS for up to 7 days, and 27 (71%) for more than a week. Protocols using 7 (*n* = 13, 34.2%) or 14 days (*n* = 12, 31.6%) of UCS were the most common. It is important to mention that some studies (*n* = 5, 13.2%) used UCS protocols of more than 15 days to explore the more severe or long-term impacts of UCS in zebrafish. As compared with the rodent literature, in which 3 to 4 weeks of UCS are necessary to observe the full behavioural phenotype, the duration of the protocols in zebrafish is remarkably shorter (predominantly 7 or 14 days). The protocols were conducted using frequently a group of up to 10 different stressors to account for unpredictability. Outcome assessment usually took place within the 24 hours following the last stress session (*n* = 31, 81.6%), with only a few studies evaluating the effects of UCS after a longer washout period (*n* = 10, 26.3%). The tests were mostly scheduled to occur at least a day from the last stressor to avoid the acute interference from the last stress session but also not too far off the end of the protocol to avoid losing the effects of UCS.

Most studies were conducted by exposing adult zebrafish to the protocol (*n* = 34, 89.4%), followed by fish in the larval (*n* = 3, 7.9%), and juvenile life stages (*n* = 1, 2.6%). Of the publications implementing the UCS protocol in early developmental stages, one of them evaluated behavioural data of the exposed animals when they were still larvae. The remaining were designed to assess the long-lasting effects of the stress, and, in this case, animals were tested more than 75 days after the protocol ended, when they were considered adults. Experiments were conducted generally with a pool of both male and female zebrafish (*n* = 21, 55.2%). In only two studies both male and female zebrafish were used and sex was analysed as a biological variable, whereas in four papers animals of only one sex were selected (*n* = 2 for male and *n* = 2 for female fish). The sex of the animals was not specified in 11 studies (28.9%). A description of the studies included in the review can be found in Table 1, and the detailed extracted information is available at https://osf.io/pbhy4. Co-authorship network analysis identified 20 clusters of researchers implementing the UCS protocol in their labs across the globe based on the studies included in this review (Fig. 2). An interactive version of the co-authorship network is available at https://tinyurl.com/2g52lbfx.

**Fig. 2.**
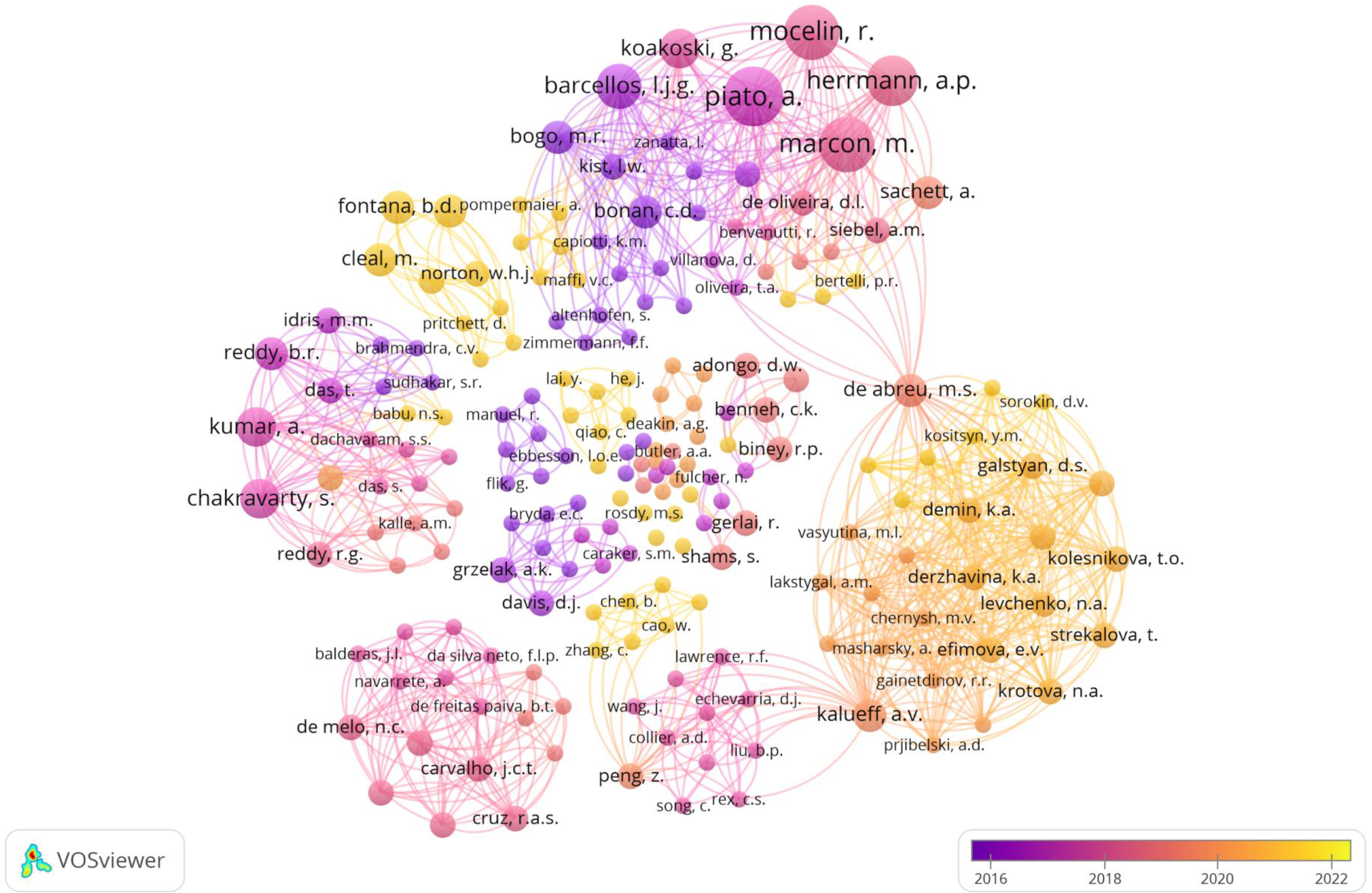
Co-authorship network analysis of researchers that authored studies implementing the unpredictable chronic stress protocol (UCS) in zebrafish. Authors are colour-coded from violet (older studies) to yellow (more recent studies) indicating the average publication year of the studies published by each researcher. The size of the circles represents the number of studies published by each author. The distance between the two circles indicates the correlations between researchers.

**Table 1.**
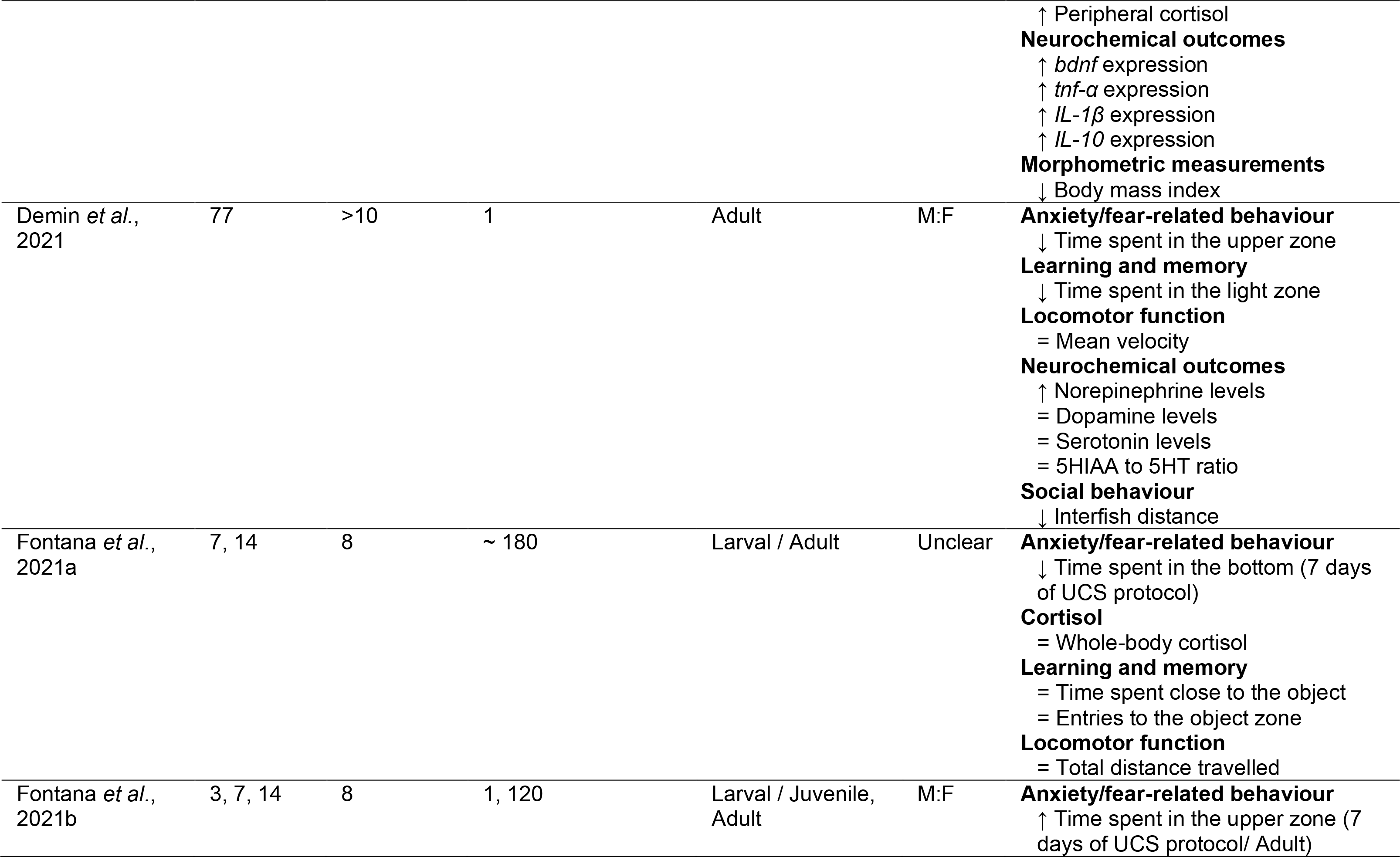
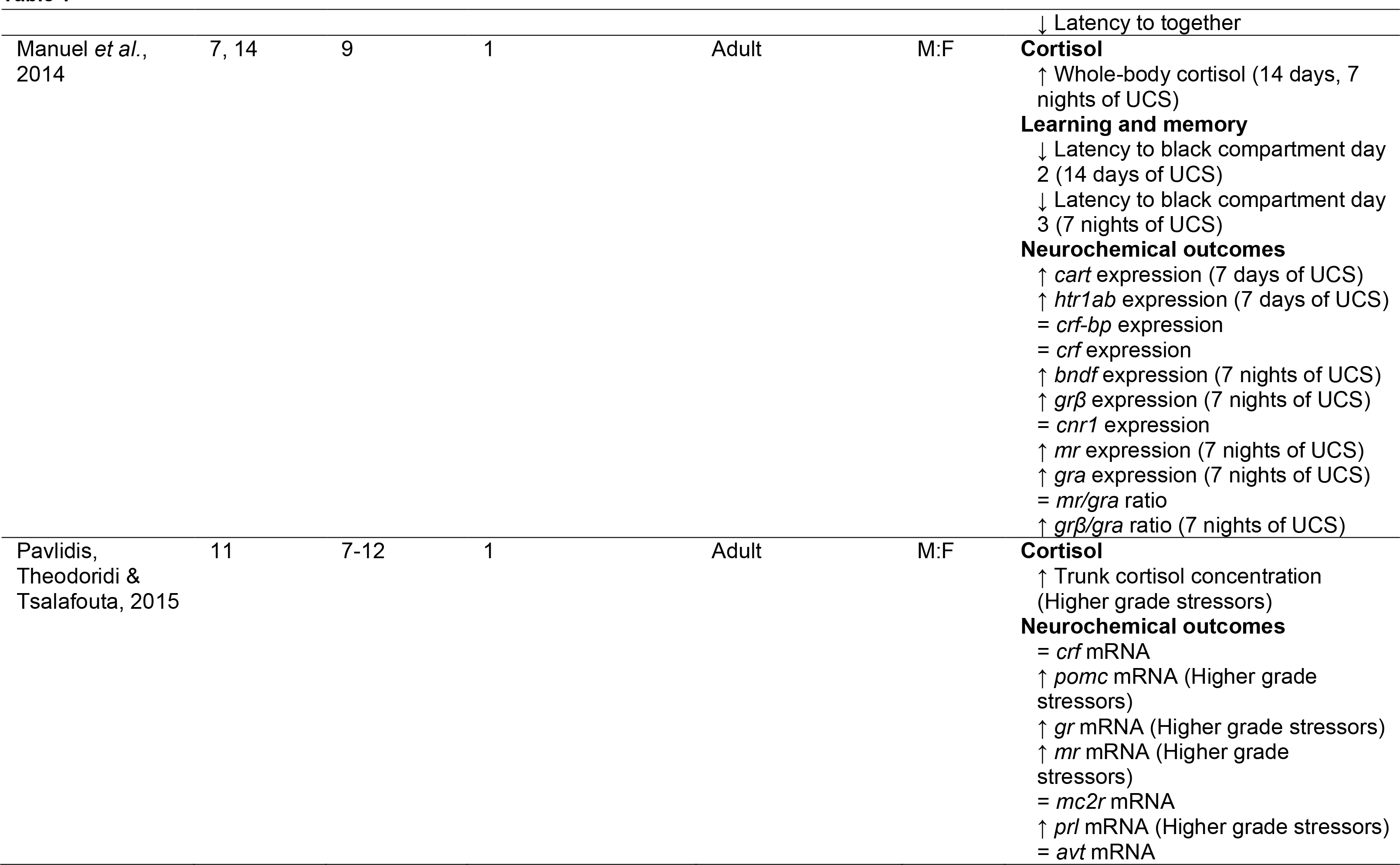
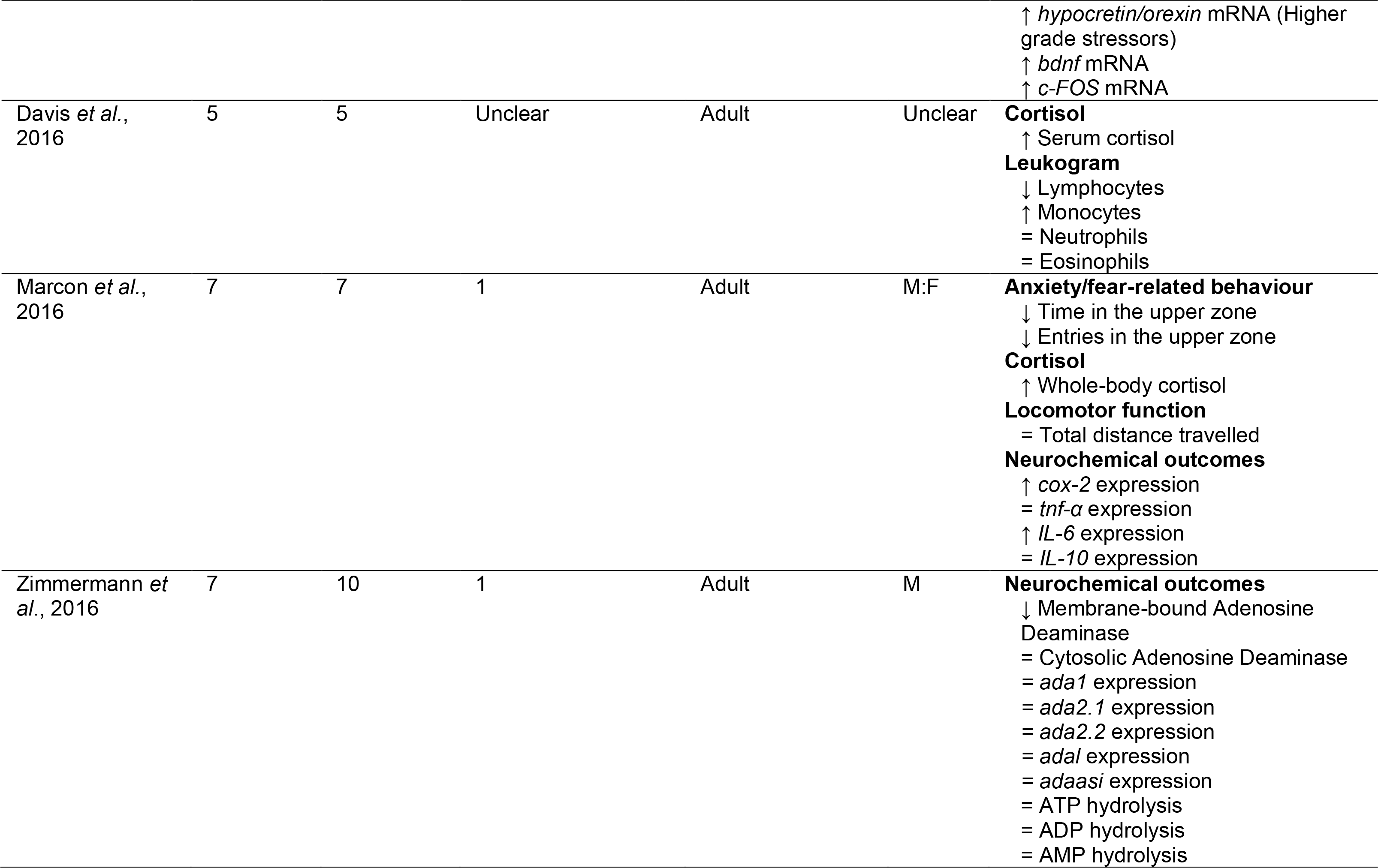
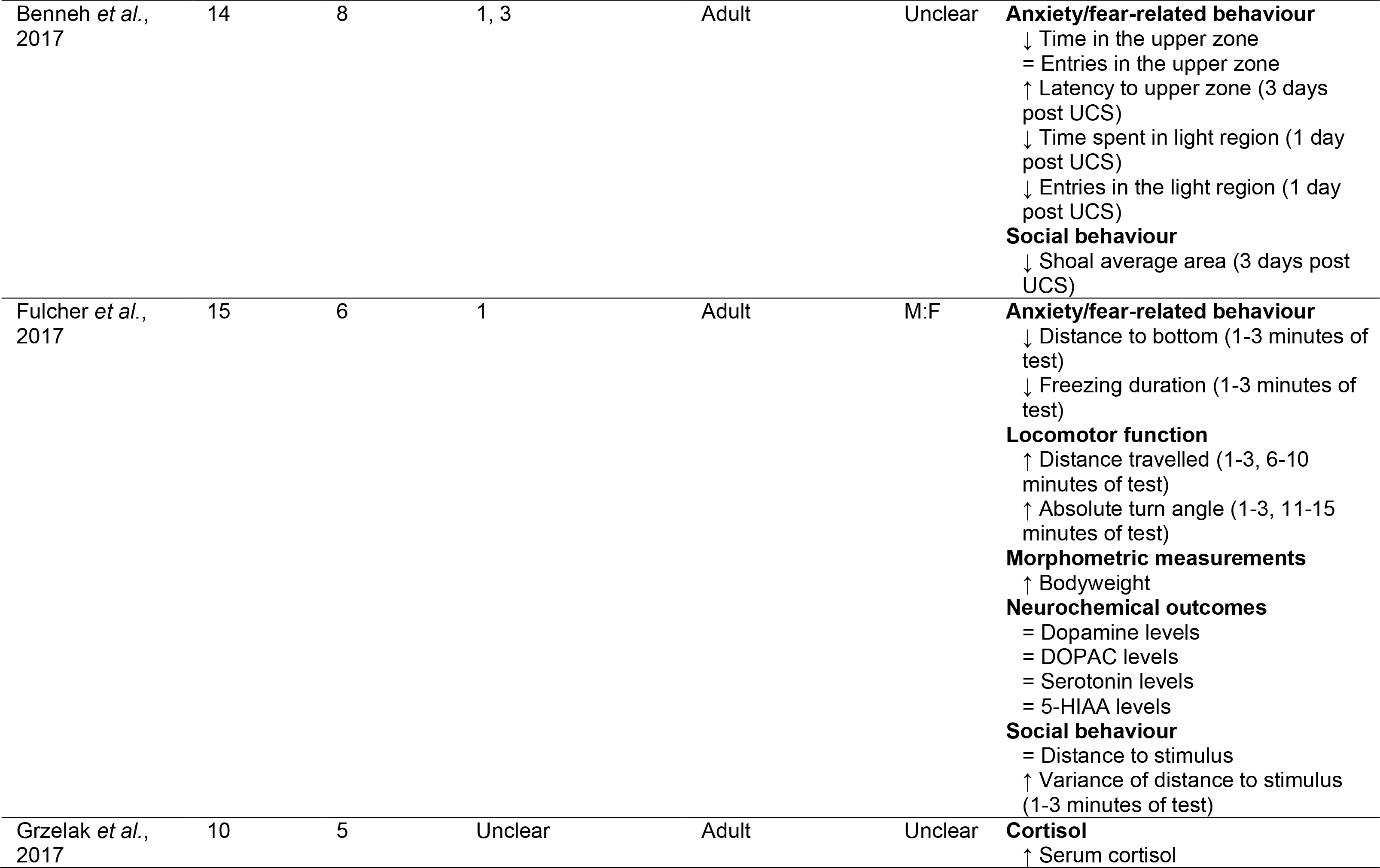
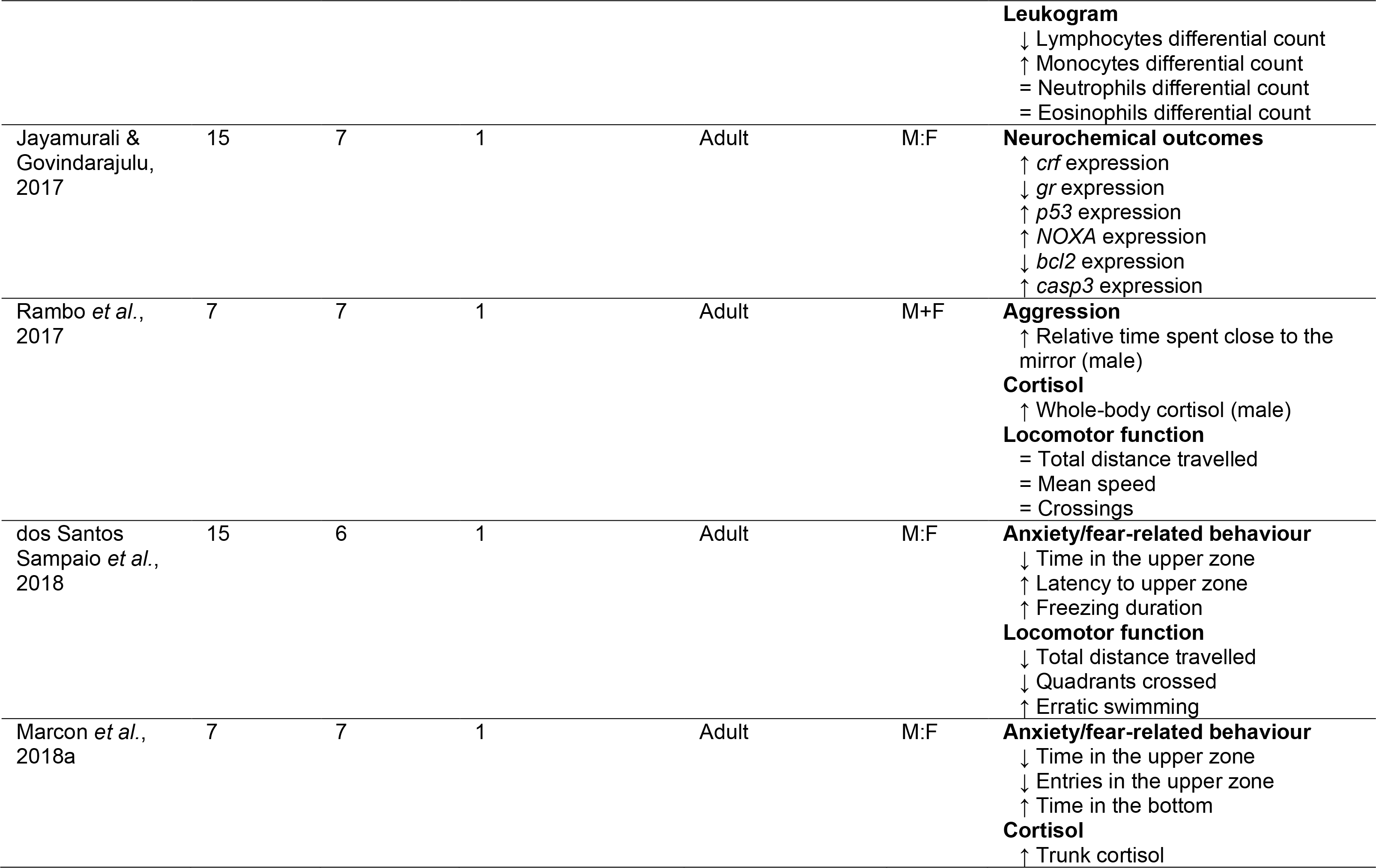
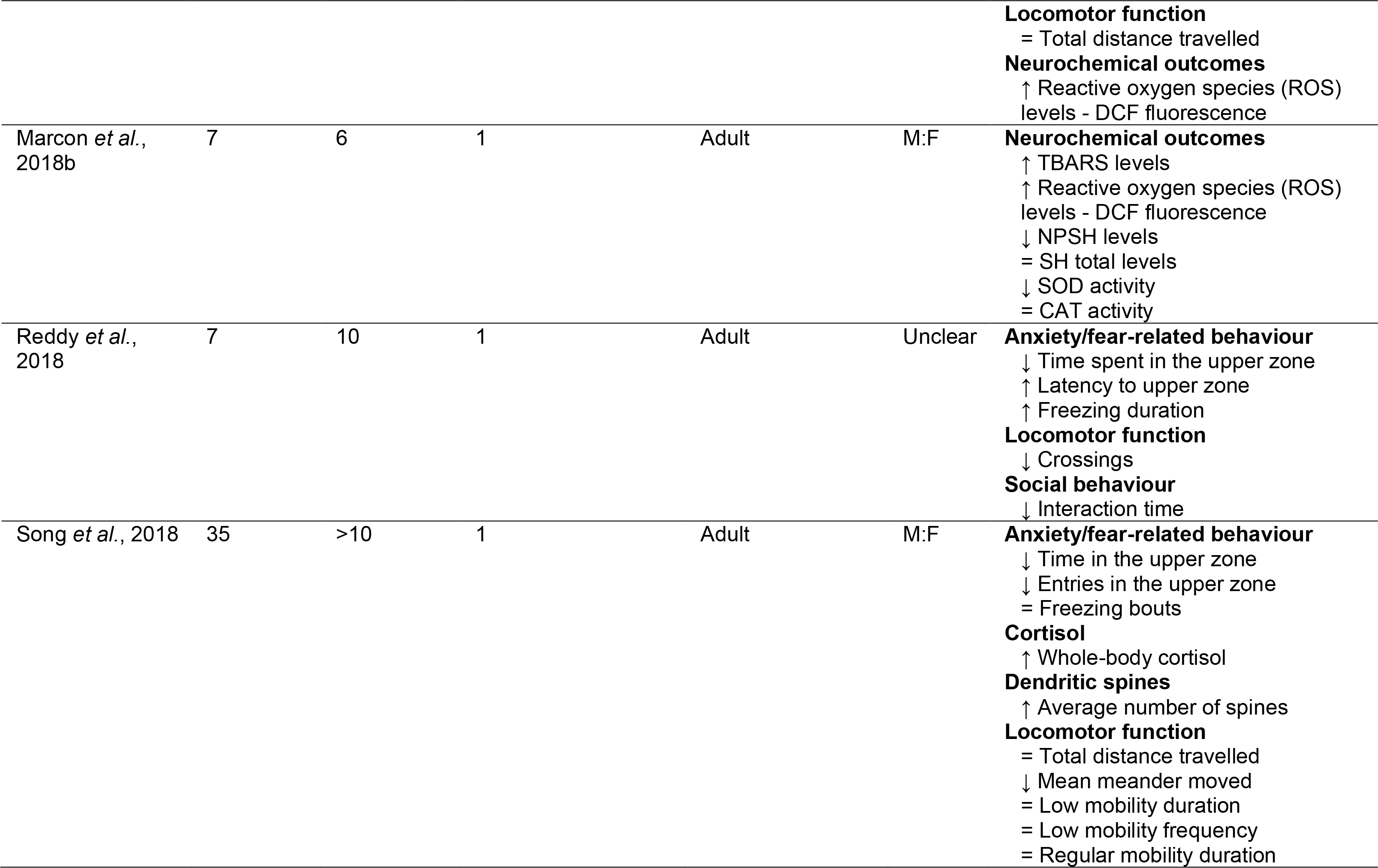
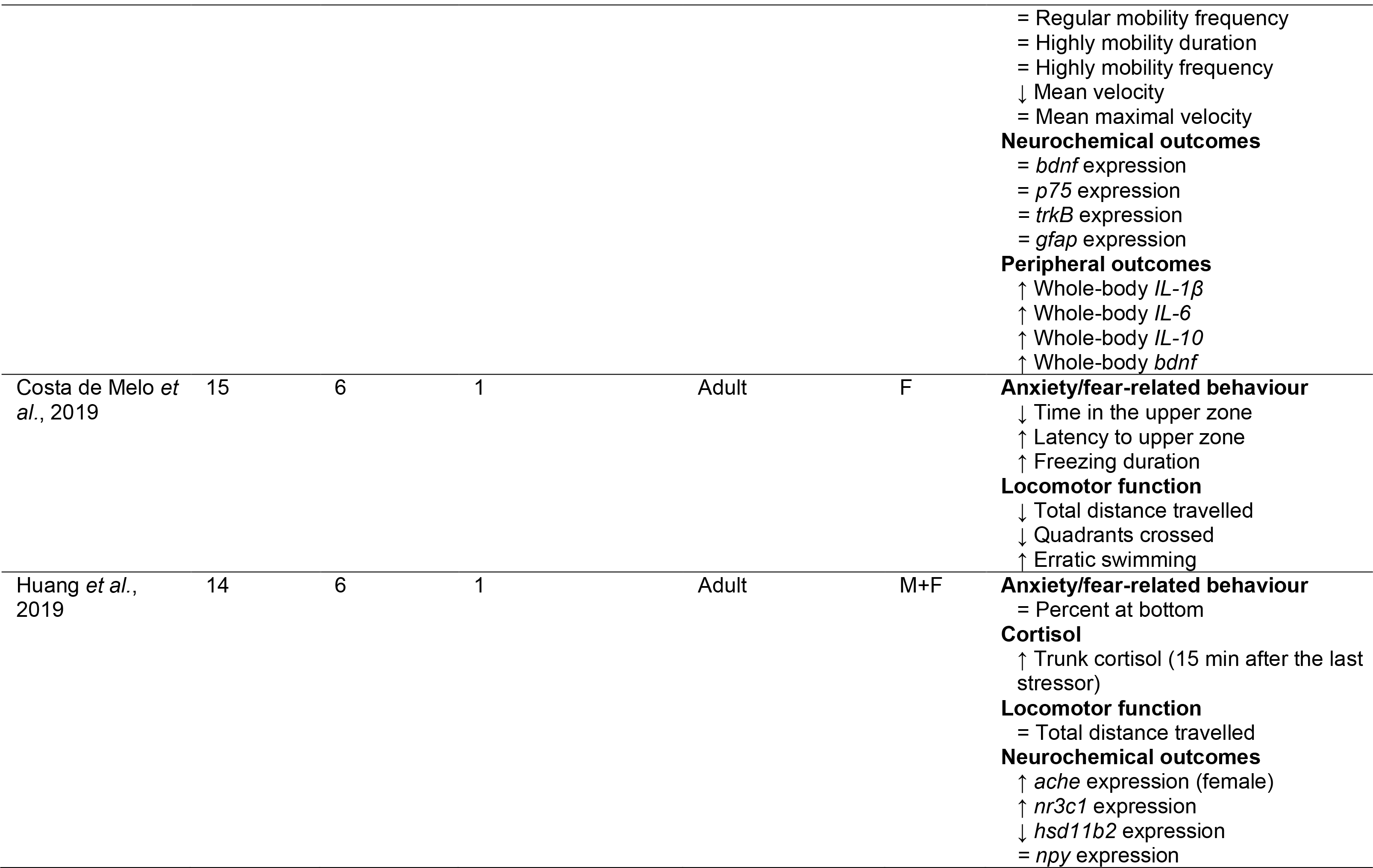
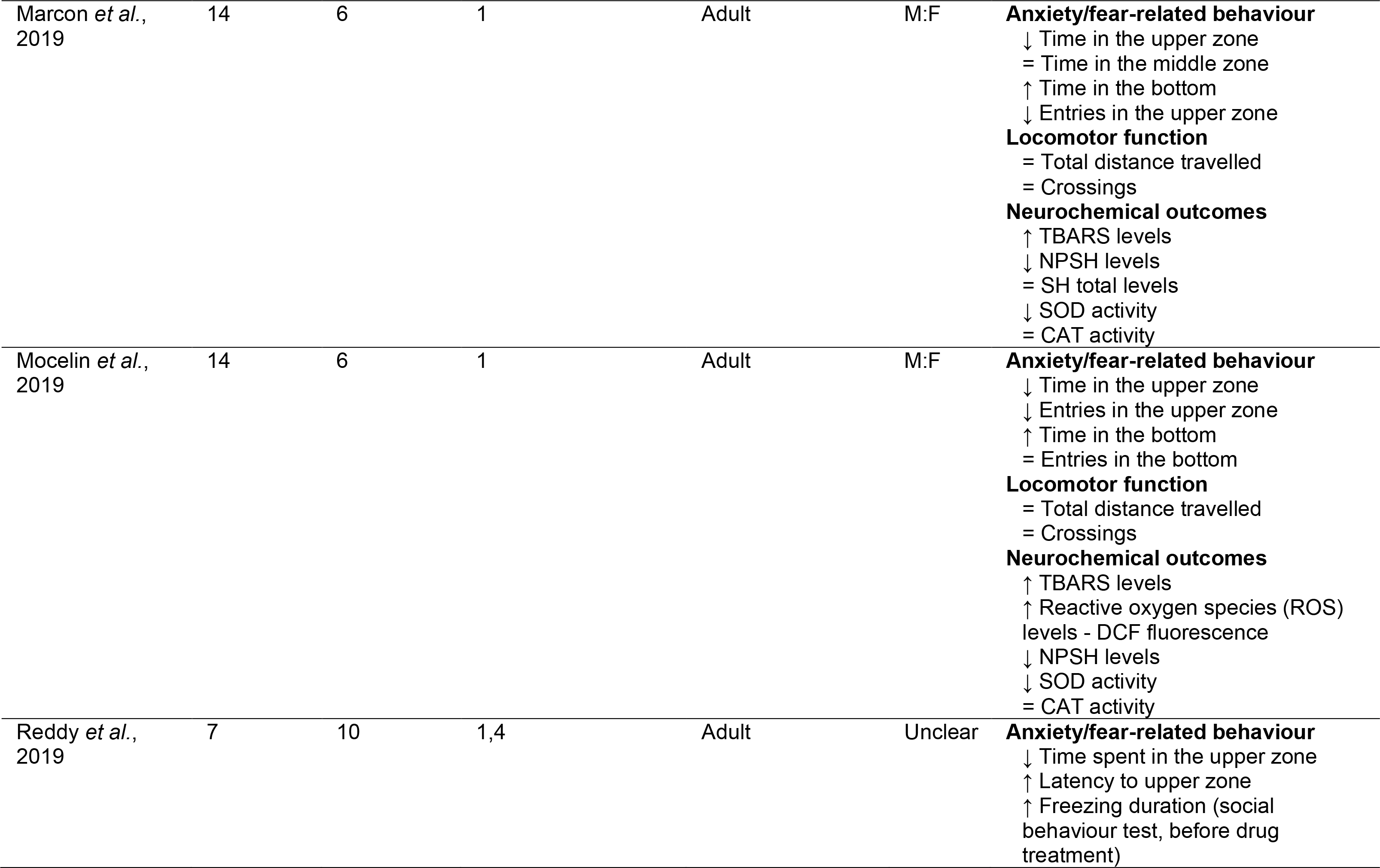
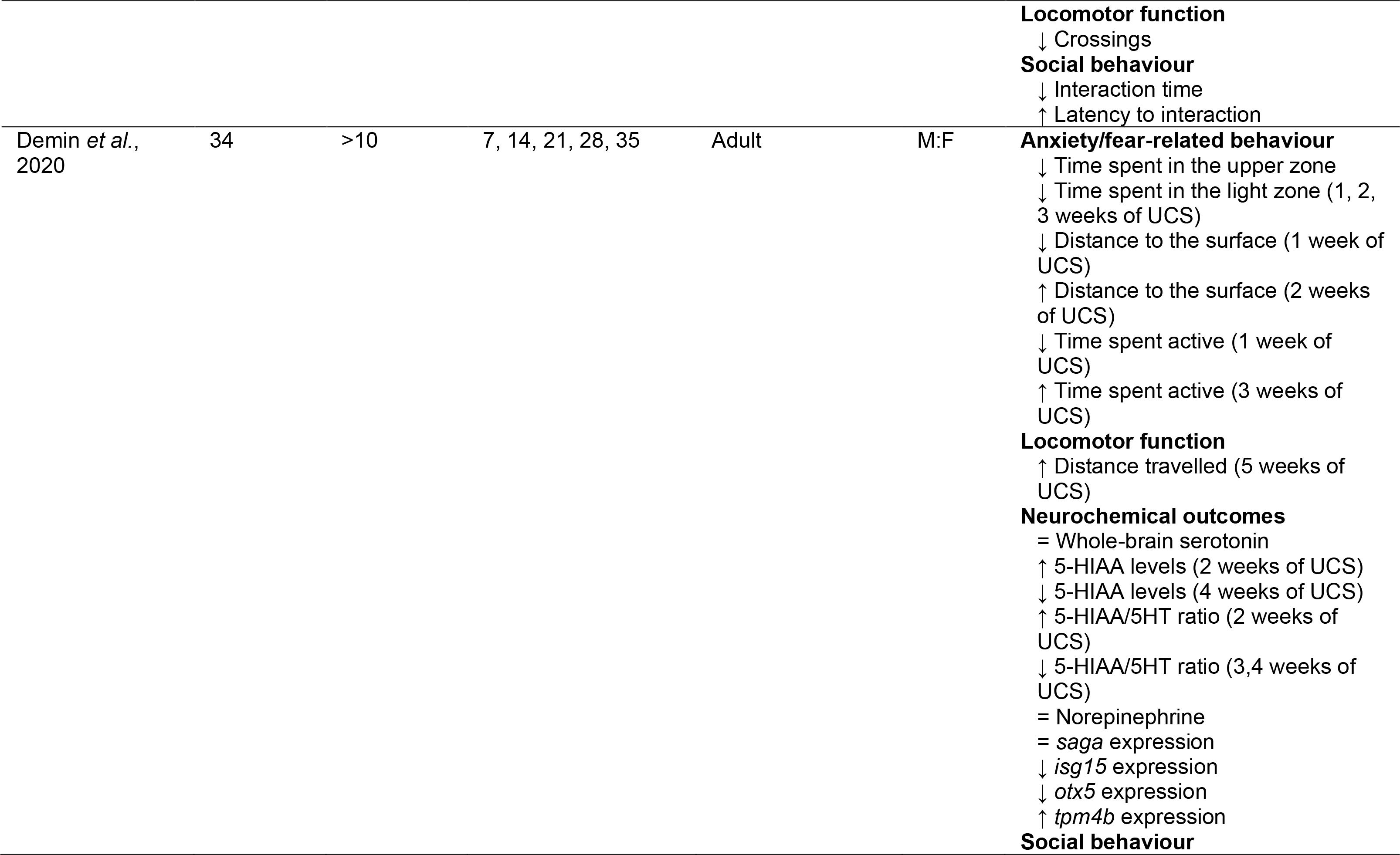
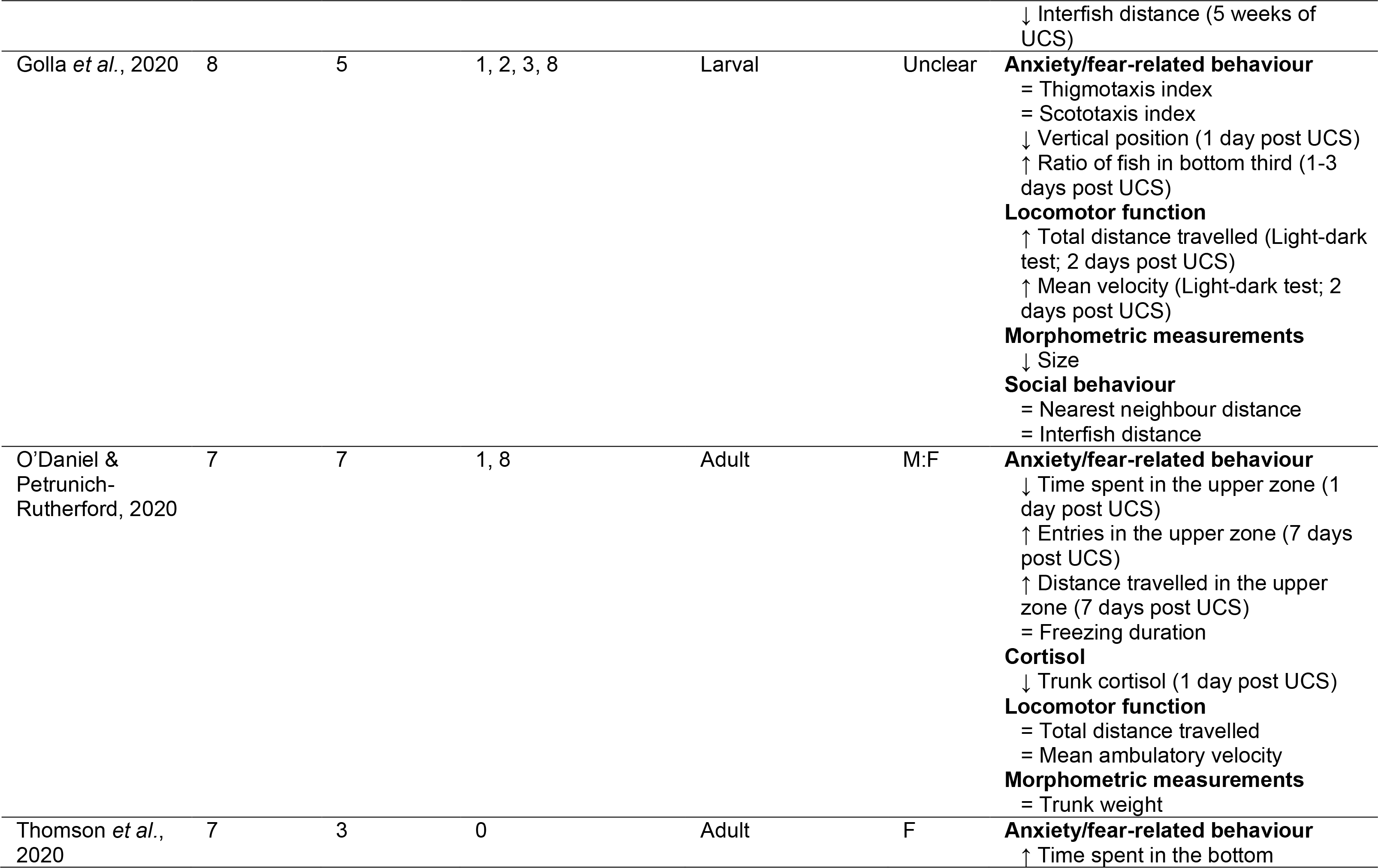
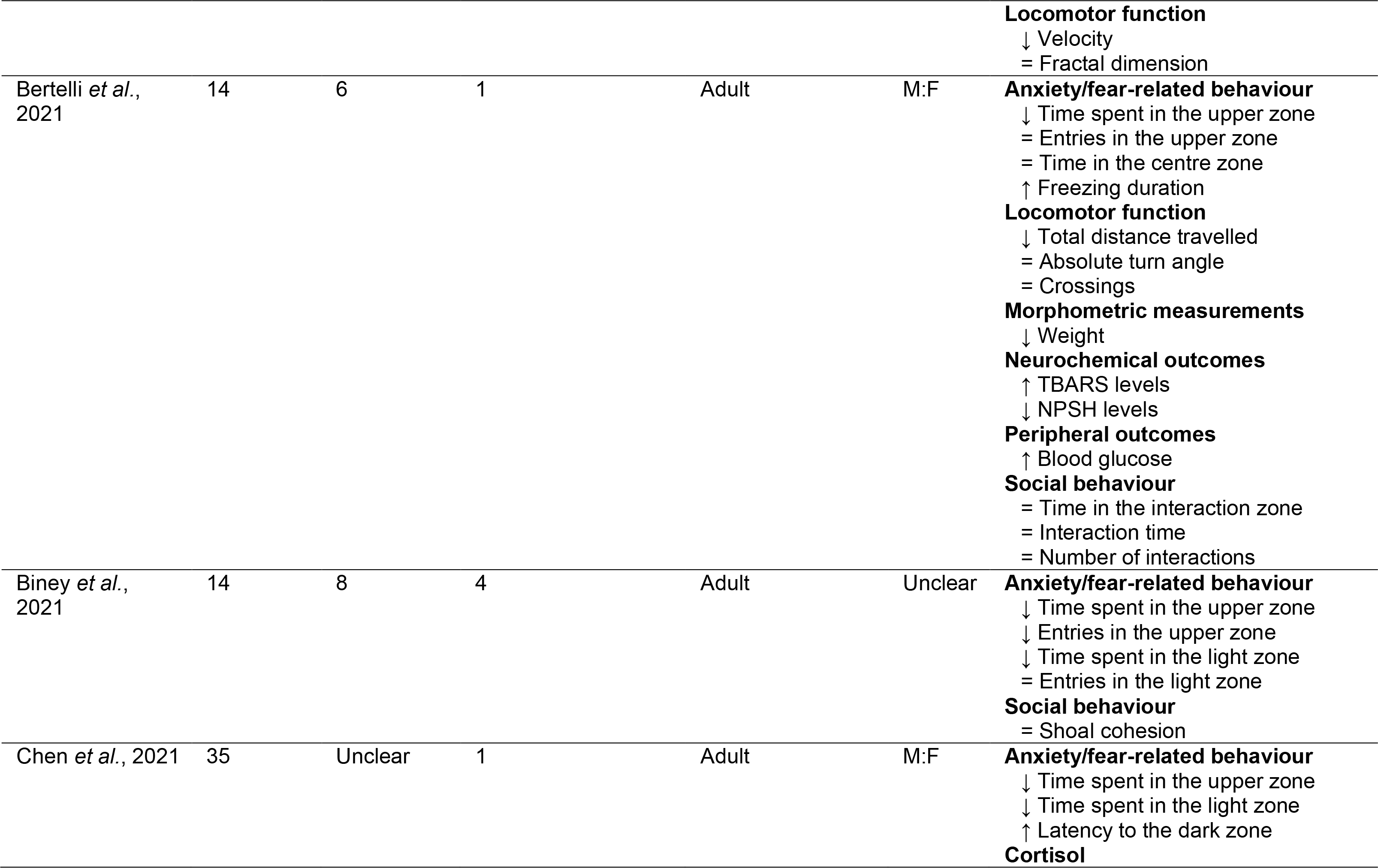
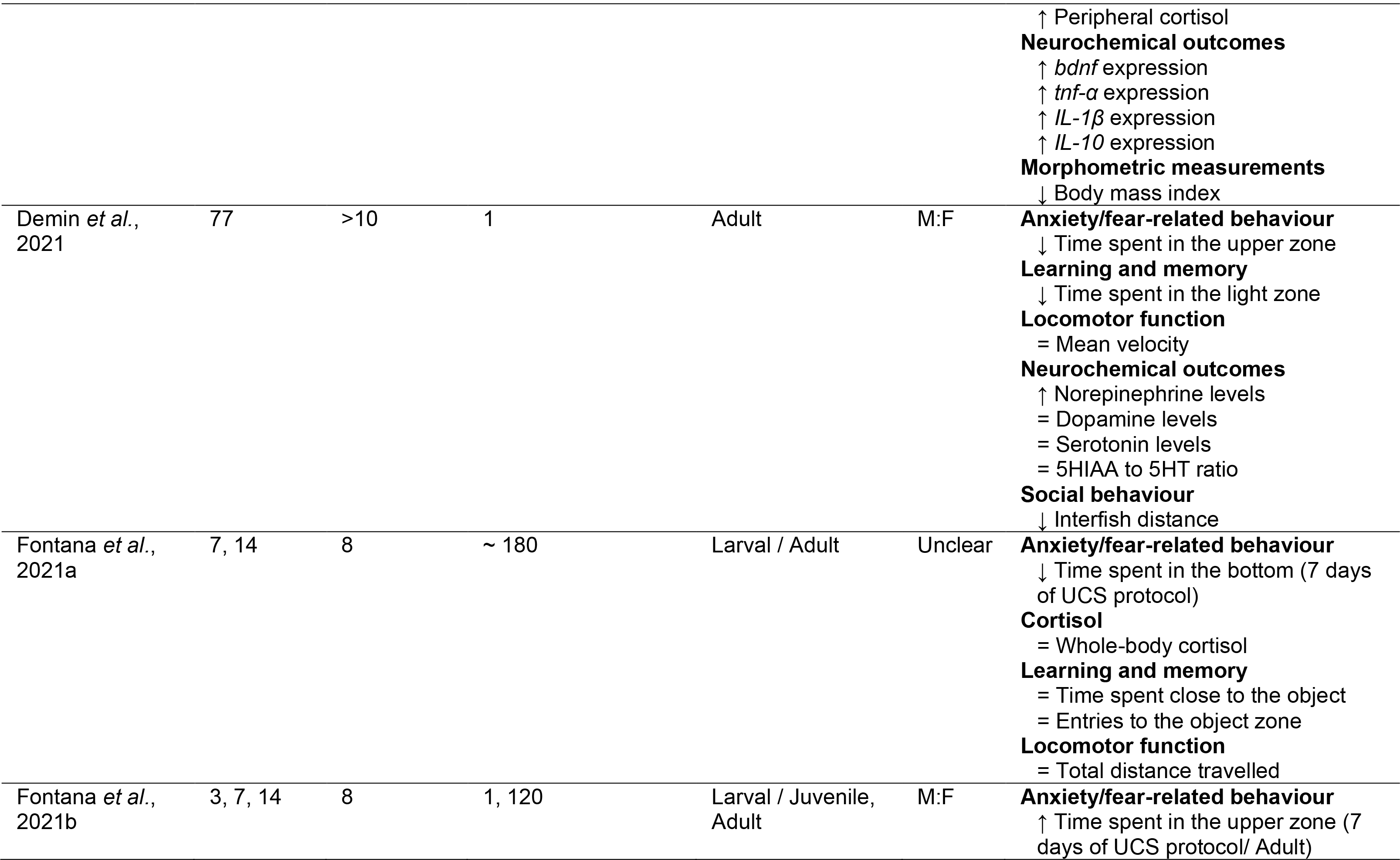
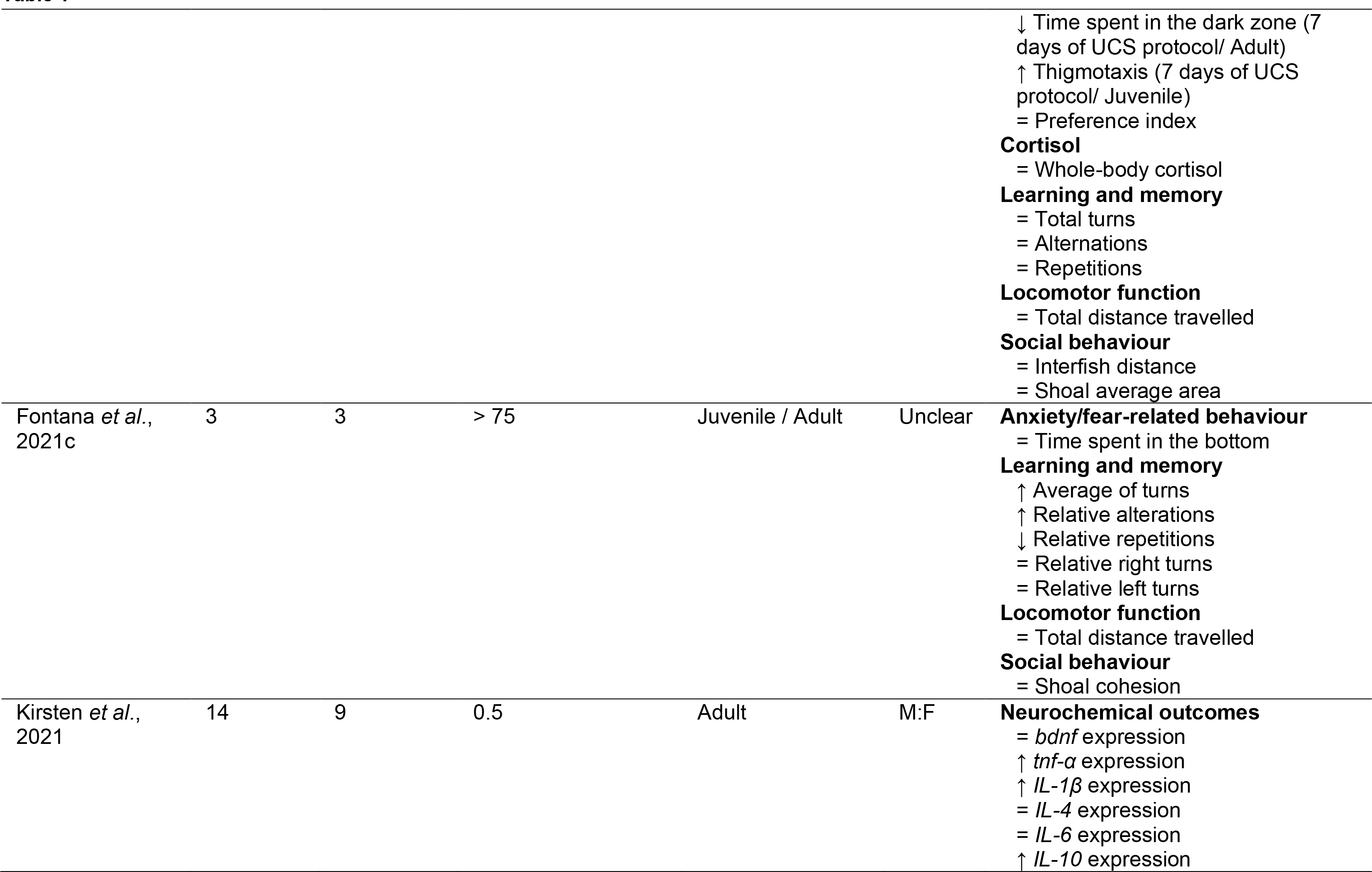
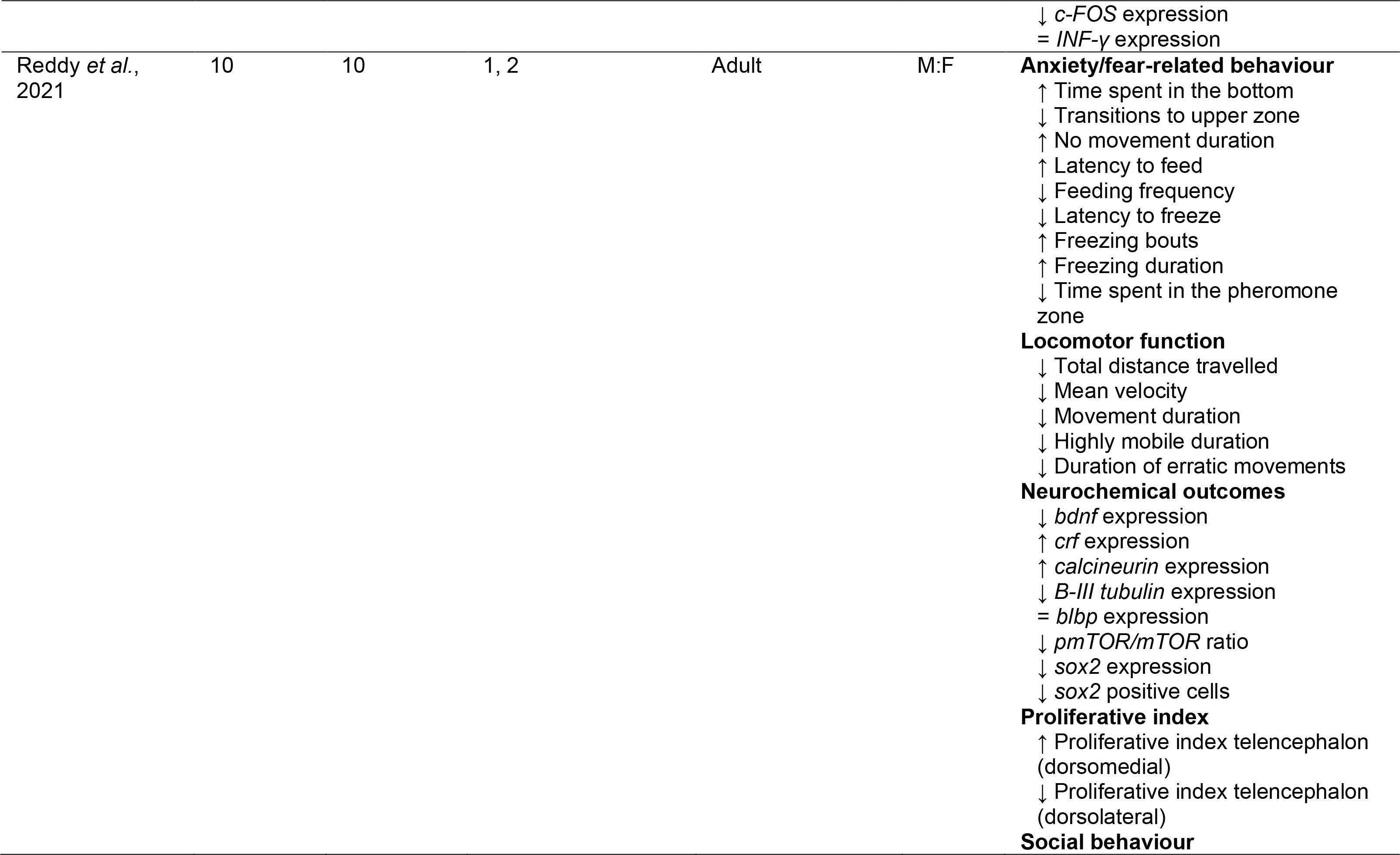
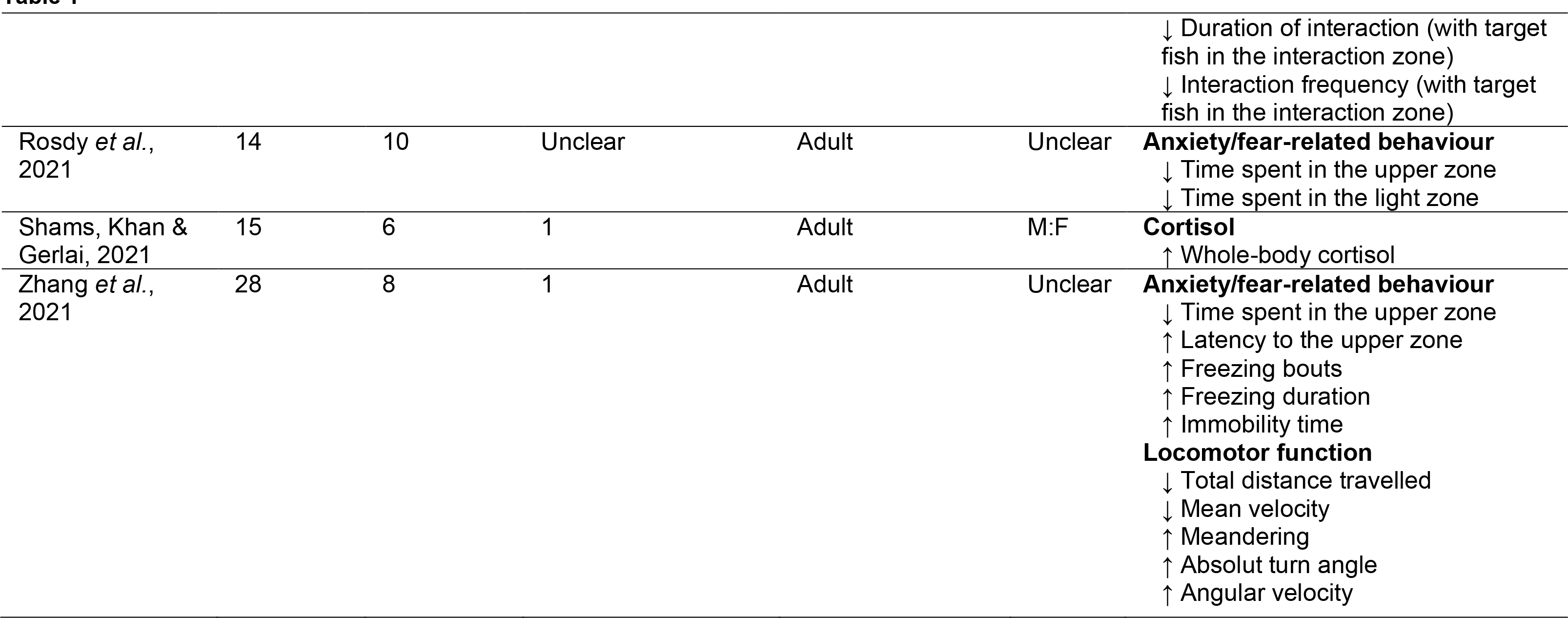
Qualitative description of studies reporting unpredictable chronic stress (UCS) protocols in research with zebrafish. The sex of the animals used was computed as: M, for male animals; F, for females; M:F, when male and female were included but tested and analyzed as a mixed group; M+F, when male and female fish were discriminated in the experiments; Unclear, for larvae and when the sex of the animals was not reported. Main findings were described as: ↑, higher when compared to the control group; ↓, lower when compared to the control group; =, no difference when compared to the control group.

### 3 Risk of bias and reporting quality

The overall risk of bias associated with the items evaluated for methodological quality was considered unclear (Fig. 3). In more than 89% of the studies included, the information given was insufficient to rule out biases arising from the allocation of animals to the experimental groups or baseline characteristics. Although being an important good research practice, random housing allocation was not reported in any publication. Bias related to blind assessment of outcomes was considered unclear in 14 studies (36.8%) and one study was deemed as having a high risk of bias for this item. Outcome data was incomplete in two studies (5.3%), and it was unclear whether data was complete in 63.2% of the assessed papers. For six studies (15.8%), cross-checking the information for outcomes measured between the methodology and the results was not possible and selective reporting was considered unclear.

**Fig. 3.**
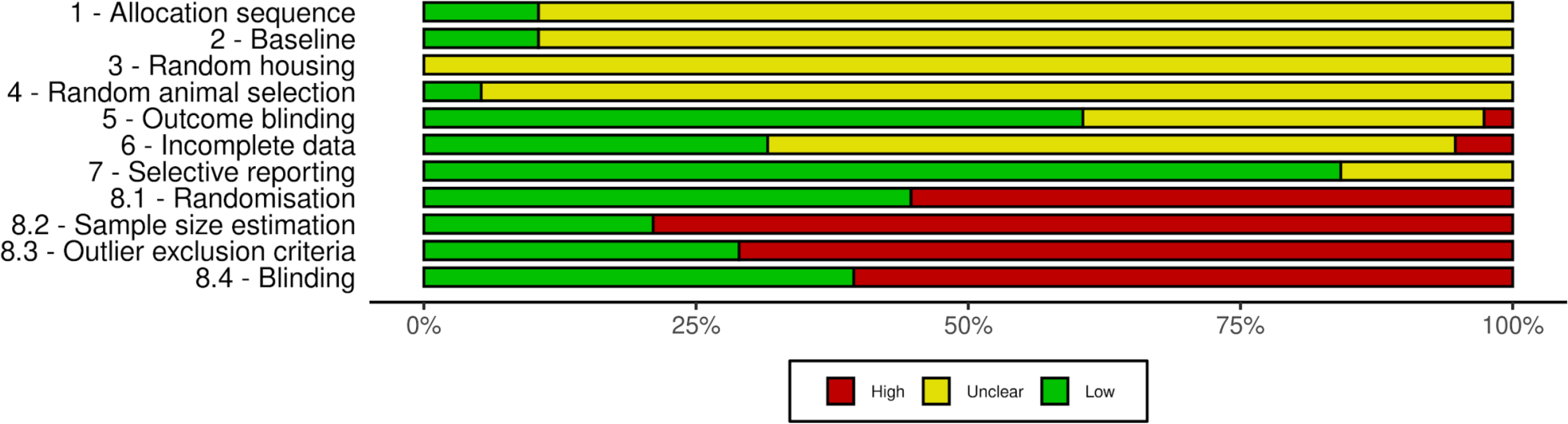
Risk of bias assessment of included studies. The risk of bias assessment was performed by two independent investigators based on the SYRCLE’s risk of bias assessment tool. Items 1 to 7 account for methodological quality and were scored as presenting a high, unclear or low risk of bias. Items 8.1 to 8.4 evaluate the reporting quality of the studies and were scored as presenting a high or low risk of bias. Classification is given as the percentage of assessed studies (*n =* 38) presenting each score.

As for the reporting quality, more than 50% of the studies failed to report any information on the items assessed. Researchers failed to describe if any randomization method was used in 21 studies (55.3%). Sample size estimation procedures were not informed in 30 papers (78.9%). Reporting quality was also considered unsatisfactory when evaluating the report of inclusion/exclusion criteria and blinding, since there were no reports of these items in 27 (71.1%) and 23 (60.5%) of the studies, respectively. Out of 418 scores given in the risk of bias assessment, there were 51 (12.2%) inconsistencies between investigators. Individualised scores for each study included are available at https://osf.io/zw6qg.

### 4 Anxiety/fear-related behaviour

The meta-analysis comprised 30 comparisons out of 22 independent studies. A total of 347 animals were used as controls and 409 composed the stressed groups. The most frequently used test to assess anxiety/fear-related behaviour in the included comparisons was the novel tank (25), followed by the open field (3), light/dark (1), and stress-induced analgesia tests (1).

The overall analysis revealed that stressed animals have higher levels of anxiety/fear-related behaviour when compared to control animals (SMD 1.09 [0.50, 1.68], *p* = 0.0007, Fig. 4). The estimated heterogeneity was high, with an I² = 81%, τ^2^ = 1.61, and a Q = 155.97 (*df* = 29, *p* < 0.01). Subgroup analysis revealed that for experiments with stress duration of up to 7 days there was no statistically significant effect on anxiety/fear-related behaviour (SMD 0.37 [-0.30, 1.04], *p* = 0.25, Fig. 4). The heterogeneity was also high for this subgroup, with an I² = 80%, a τ^2^ = 0.79, and a Q = 49.83 (*p* < 0.01). For experiments with a UCS regimen of more than 7 days, it is possible to observe a significant effect of the stress on increasing anxiety-like behaviour (SMD 1.58 [0.73, 2.43], *p* < 0.01, Fig. 4). The heterogeneity remained high when analysing this subgroup, resulting in an I² = 80%, a τ^2^ = 2.06, and a Q = 88.60 (*p* < 0.01). The difference between subgroups was also significant (*p* = 0.02) suggesting that the duration of the UCS protocol modifies the effect of the stress on anxiety/fear-related behaviour of zebrafish.

**Fig. 4.**
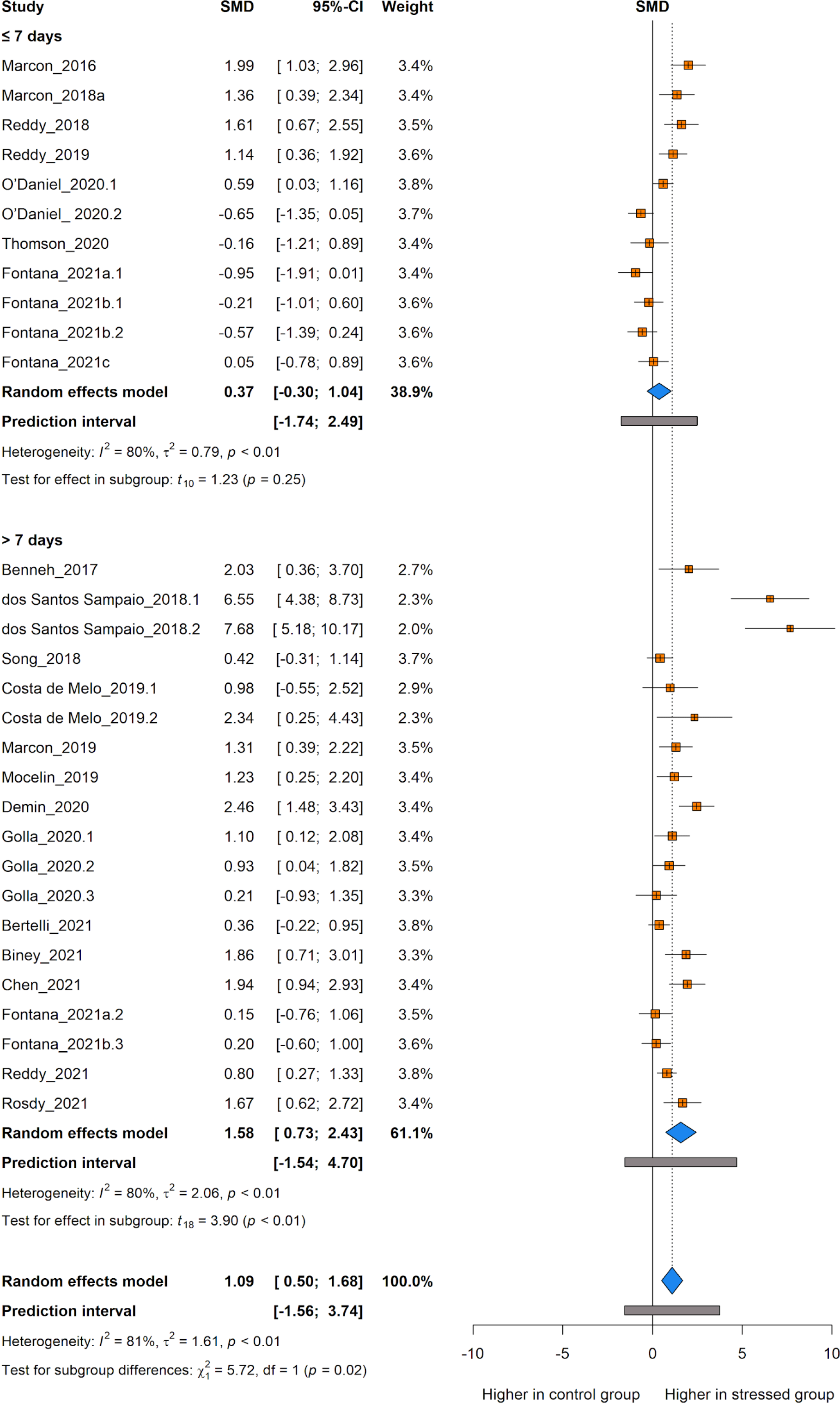
The effect of unpredictable chronic stress (UCS) protocol on anxiety/fear- related behaviour of zebrafish. Subgroup analyses were based on the duration of the stress protocol (either ≤ 7 days or > 7 days of stress). Data are presented as Hedges’ G standardised mean differences (SMD) and 95% confidence intervals.

### 5 Locomotor function

The meta-analysis comprised 28 comparisons out of 21 independent studies. A total of 454 animals were used as controls and 510 composed the stressed groups. The most frequently used test to assess locomotor function in the included studies was the novel tank (21), followed by the open field (4), mirror-induced aggression (2), and stress-induced analgesia tests (1).

The overall analysis showed that stressed animals show lower levels of mobility when compared to control animals (SMD -0.56 [-1.02, -0.10], *p* = 0.0180, Fig. 5). The estimated heterogeneity was considered high, with an I² = 83%, τ^2^ = 1.01, and a Q = 156.83 (*df* = 27, *p* < 0.01). When analysing separately experiments conducted with a UCS protocol of up to 7 days, there was no statistically significant effect of the stress on locomotor function (SMD -0.21 [-0.74, 0.33], *p* = 0.42, Fig. 5). The heterogeneity was also high for this subgroup, with an I² = 76%, a τ^2^ = 0.59, and a Q = 49.51 (*p* < 0.01). As for experiments conducted with a UCS regimen of more than 7 days, it is possible to observe a significant difference in locomotor function between stressed and control groups, evidencing lower mobility in stressed animals (SMD -0.93 [-1.69, -0.16], *p* = 0.02, Fig. 5). The heterogeneity remained high when analysing this subgroup, resulting in an I² = 86%, a τ^2^ = 1.43, and a Q = 101.67 (*p* < 0.01). The difference between subgroups was also significant (*p* = 0.10) suggesting that the duration of the UCS protocol modifies the effect of the stress on the locomotor function of zebrafish.

**Fig. 5.**
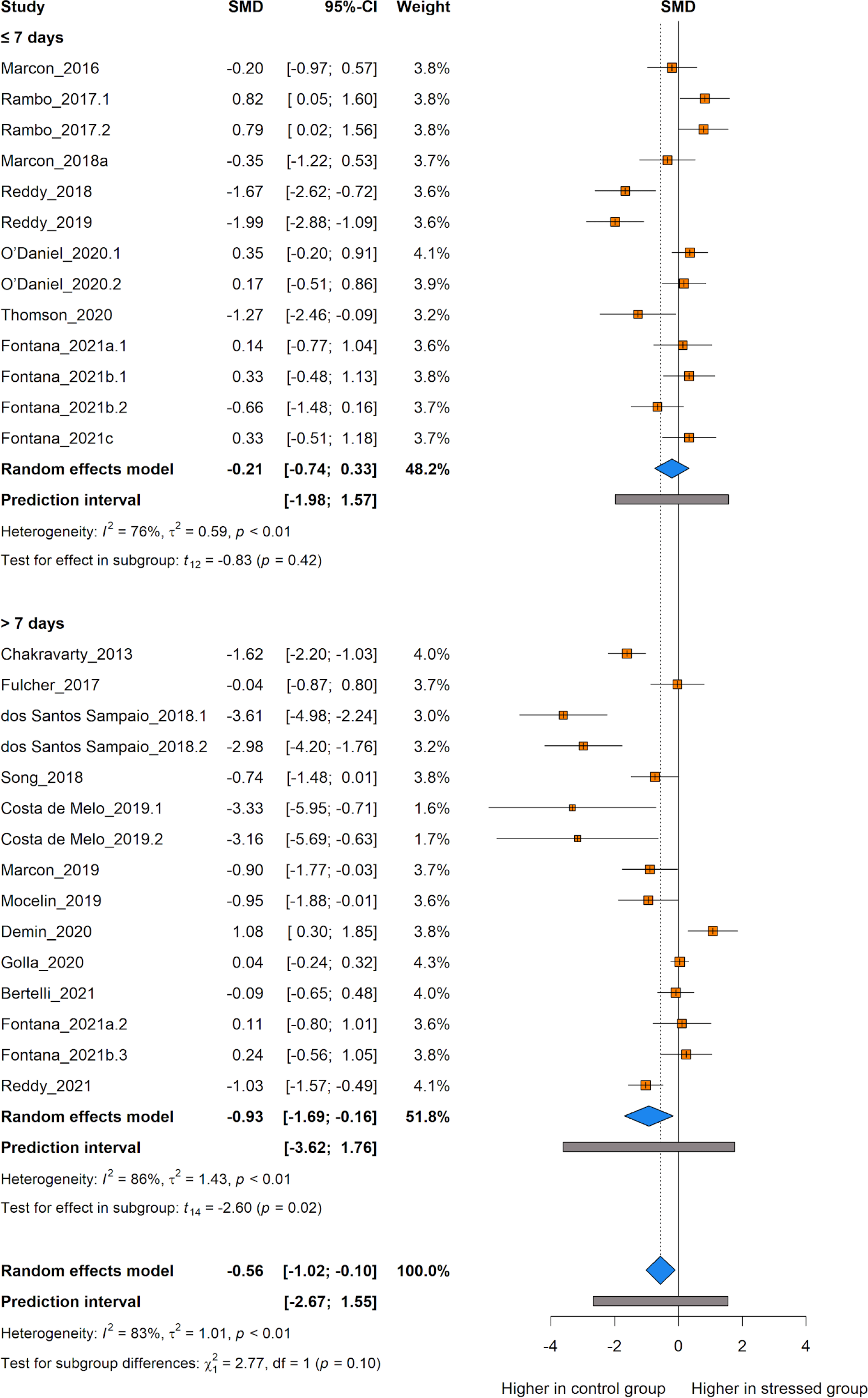
The effect of unpredictable chronic stress (UCS) protocol on the locomotor function of zebrafish. Subgroup analyses were based on the duration of the stress protocol (either ≤ 7 days or > 7 days of stress). Data are presented as Hedges’ G standardised mean differences (SMD) and 95% confidence intervals.

### 6 Social behaviour

The meta-analysis comprised 14 comparisons out of 11 independent studies. A total of 172 animals were used as controls and 190 composed the stressed groups. The most frequently used test to assess social behaviour in the included studies was the shoaling response test (8), followed by social interaction (4), and novel tank tests (2). The overall analysis showed no significant effects of the UCS protocol on social behaviour (SMD -0.30 [-0.77, 0.17], *p* = 0.1849, Fig. 6). The estimated heterogeneity was considered moderate, with an I² = 74%, a τ^2^ = 0.47, and a Q = 50.86 (*df* = 13, *p* < 0.01). There were no sufficient studies to perform a subgroup analysis.

**Fig. 6.**
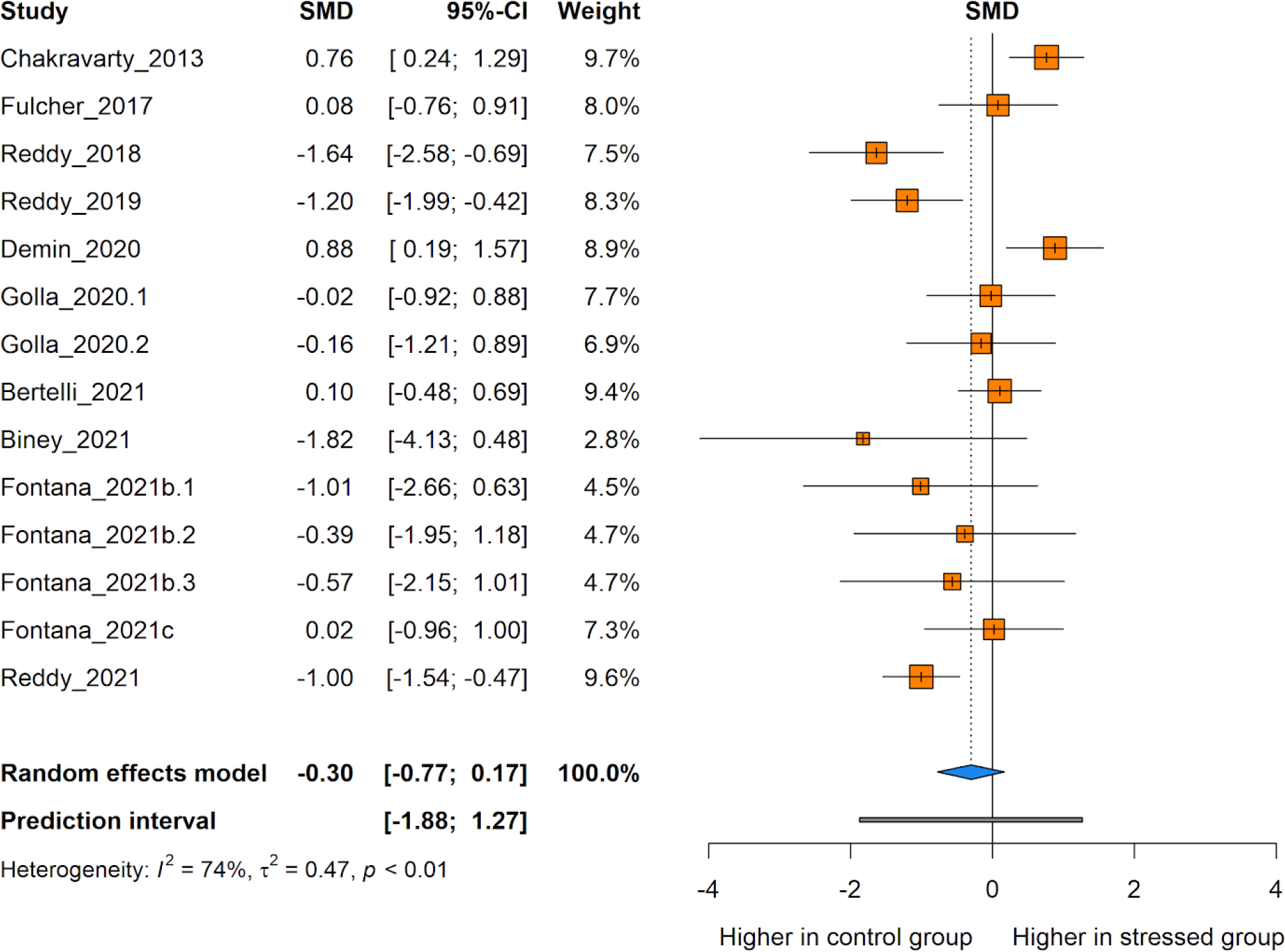
The effect of unpredictable chronic stress (UCS) protocol on the social behaviour of zebrafish. Data are presented as Hedges’ G standardised mean differences (SMD) and 95% confidence intervals.

### 7 Cortisol levels

The meta-analysis comprised 22 comparisons out of 13 independent studies. A total of 150 animals were used as controls and 223 composed the stressed groups. Whole-body cortisol levels were measured in most studies (15), followed by trunk (5), and serum cortisol measurements (2).

The overall analysis showed that stressed animals have higher levels of cortisol when compared to control animals (SMD 0.66 [0.06, 1.25], *p* = 0.0320, Fig. 7). The estimated heterogeneity was considered moderate, with an I² = 75%, a τ^2^ = 1.01 and a Q = 84.98 (*df* = 21, *p* < 0.01). When analysing separately experiments conducted with a UCS regimen of up to 7 days, there was no statistically significant effect of the stress on cortisol levels (SMD 0.73 [-0.27, 1.74], *p* = 0.14, Fig. 7). The heterogeneity was high for this subgroup, with an I² = 83%, a τ^2^ = 2.00, and a Q = 77.02 (*p* < 0.01).

**Fig. 7.**
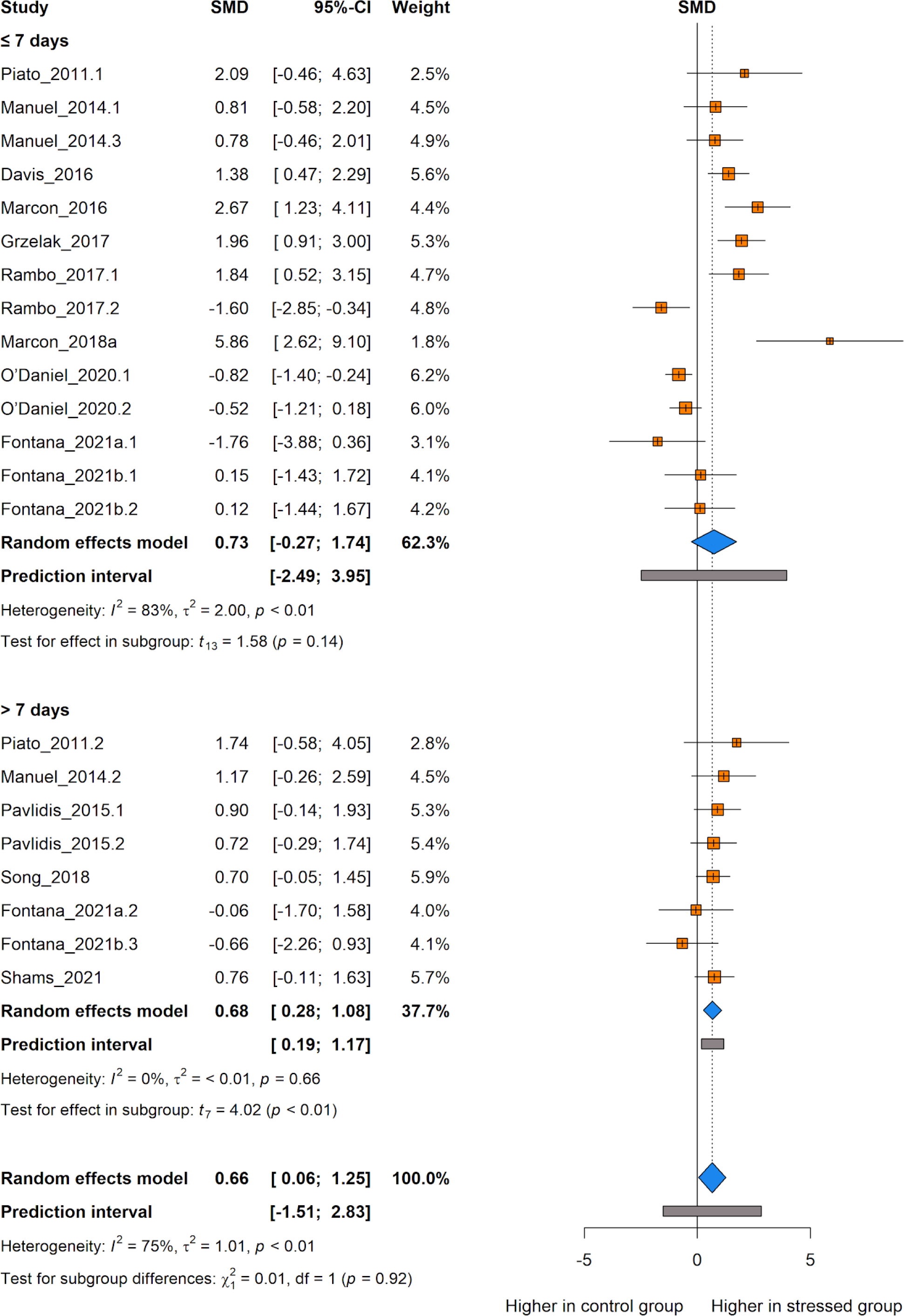
The effect of unpredictable chronic stress (UCS) protocol on cortisol levels in zebrafish. Subgroup analyses were based on the duration of the stress protocol (either ≤ 7 days or > 7 days of stress). Data are presented as Hedges’ G standardised mean differences (SMD) and 95% confidence intervals.

As for experiments conducted with a UCS protocol of more than 7 days, it is possible to observe a significant effect of the stress on increasing cortisol levels (SMD 0.68 [0.28, 1.08], *p* < 0.01, Fig. 7). The heterogeneity significantly decreased when analysing this subgroup, resulting in an I² = 0%, a τ^2^ < 0.01 and a Q = 4.98 (*p* = 0.66).

The difference between subgroups was not statistically significant (*p* = 0.92), suggesting that the duration of the UCS protocol does not modify the effect of the stress on the cortisol levels in zebrafish.

### 8 Publication bias

Visual inspection of funnel plots demonstrated a substantial asymmetrical distribution of the studies within some domains of interest (Fig. 8). The scattered plot does not show the expected funnel-shaped distribution of experiments for anxiety/fear-related behaviour (Fig. 8A), locomotor function (Fig. 8B), social behaviour (Fig. 8C), and cortisol levels (Fig. 8D). This could be attributed to sample heterogeneity, as the protocols, tests, and measured variables differ significantly among selected studies. Trim and fill analysis for anxiety/fear-related behaviour imputed 8 studies to the meta-analysis, and the overall effect of the stress was not significant for this outcome (SMD 0.50 [-0.23, 1.24], *p* = 0.1746, Fig. 8A). For locomotor function, 5 studies were imputed, and the overall effect of the stress was not significant (SMD -0.20 [-0.77, 0.36], *p* = 0.4677, Fig. 8B). For social behaviour, 6 studies were imputed, and the overall effect of the stress remained not significant (SMD 0.28 [-0.31, 0.86], *p* = 0.3338, Fig. 8C). For cortisol levels, 5 studies were imputed, and the overall effect of the stress was once again not significant (SMD 0.20 [-0.51, 0.91], *p* = 0.5683, Fig. 8D).

**Fig. 8.**
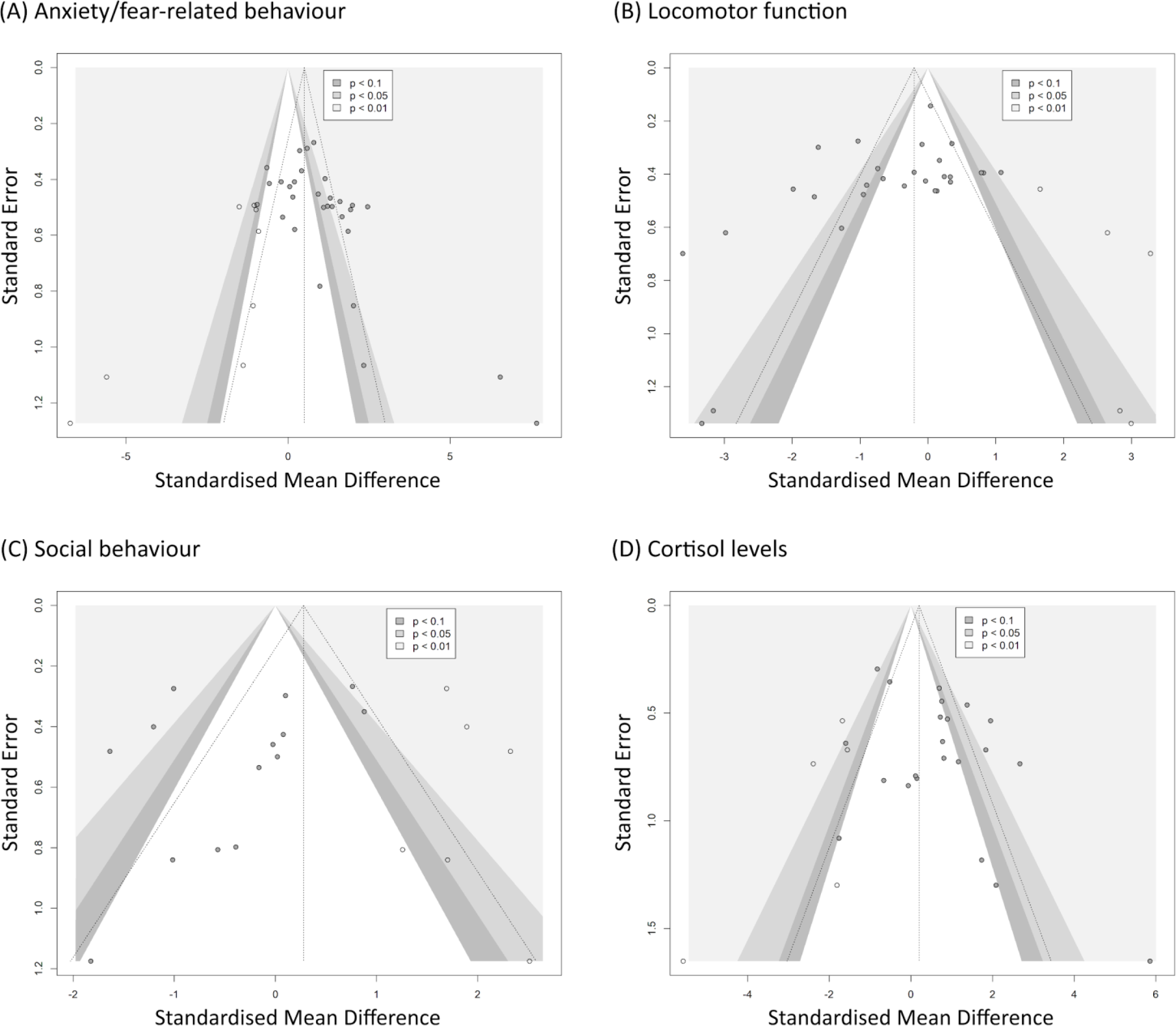
Funnel plots including studies analysed within each domain of interest: (A) anxiety/fear-related behaviour, (B) locomotor function, (C) social behaviour, and (D) cortisol levels. Each grey circle represents a single comparison. Hollow circles represent imputed studies in the trim and fill analysis. The vertical line represents the overall effect size and the triangular region represents the 95% confidence interval. Shaded areas represent the interval for statistically significant effects.

Egger’s regression test indicated publication bias for most domains tested (Table 2): anxiety/fear-related behaviour (*p* = 0.001), locomotor function (*p* = 0.0277), and cortisol levels (*p* = 0.0566). All tests suggest a possible overestimation of the effects of UCS based on published data. For social behaviour, on the other hand, the regression test was not statistically significant (*p* = 0.2507), but this result should be interpreted with caution as only a few studies reported such outcome and the statistical power of the Egger’s test heavily depends on the number of experiments included in the analysis.

**Table 2.** Regression test for Funnel plot asymmetry (“Egger’s test”). A p-value < 0.1 was considered significant for publication bias.

| Domain | Intercept | Standard Error | t | $p$-value |
| --- | --- | --- | --- | --- |
| Anxiety/fear-related behaviour | -1.0856 | 0.5182 | 3.68 | 0.001 |
| Locomotor function | 0.5310 | 0.3866 | -2.33 | 0.0277 |
| Social Behaviour | 0.4880 | 0.5840 | -1.21 | 0.2507 |
| Cortisol levels | -0.7010 | 0.5919 | 2.02 | 0.0566 |

### 9 Sensitivity analysis

The sensitivity analyses for studies presenting a significant risk of bias skewed the main effect of the domains tested (Fig. 9). After excluding studies with a high risk of bias, no significant effects of UCS on anxiety/fear-related behaviour (SMD 1.01 [-0.19, 2.22], Fig. 9A) and locomotor function (SMD -0.42 [-1.14, 0.30], Fig. 9B) were observed. For social behaviour, the overall interpretation remained the same, with no significant effects of the intervention on this behaviour (SMD 0.07 [-0.46, 0.60], Fig. 9C). For cortisol levels, on the other hand, by excluding studies associated with a high risk of bias the direction of the effect was reversed, as the meta-analysis evidenced higher levels of cortisol in the control animals when compared to the stressed groups (SMD -0.60 [-0.98, -0.21], Fig. 9D). Although it is an interesting result, the meta-analysis conducted with studies presenting a low risk of bias for the cortisol levels is based only on the results of three individual studies with different experiments, which hinders the extrapolation of this result. In the sensitivity analyses following the jackknife method, no study was shown to be skewing the overall result of the meta-analyses as the interpretation of the summary results was altered by omitting one study at a time (Suppl. Fig. 1). An exception to this was the sensitivity analysis for cortisol: in the absence of four of the studies the overall effect size is not statistically significant. This, however, is not surprising considering that the lower limit of the confidence interval in the original meta-analysis was already close to zero. Furthermore, heterogeneity is not much altered in any of the leave-one-out simulations as compared to the original result.

**Fig. 9.**
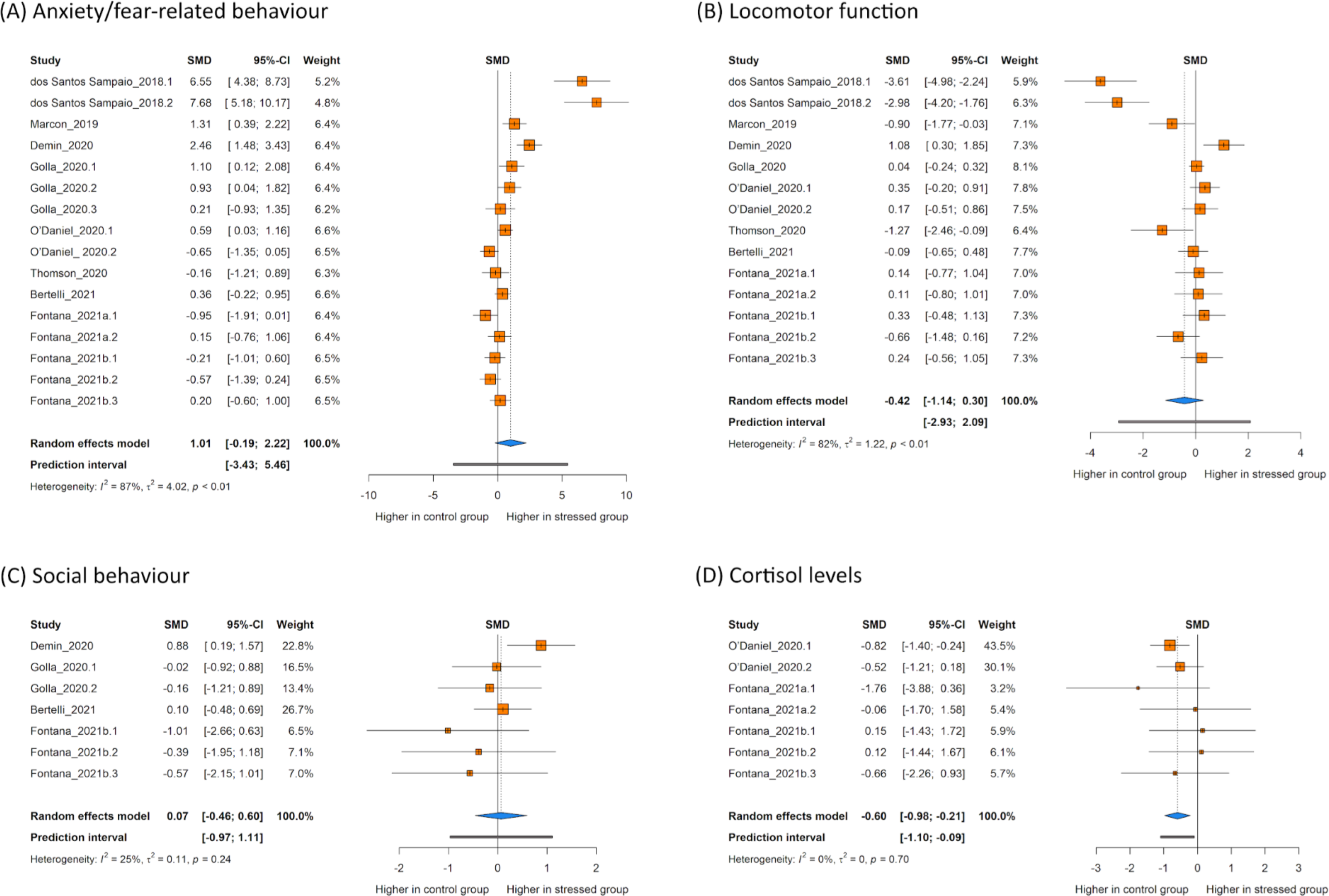
Sensitivity analyses for studies with a high risk of bias. The analyses were conducted by excluding studies presenting a significant risk of bias, defined as either a high risk of bias in one of the main items evaluating methodological quality in the risk of bias assessment (items 1 to 7), or an unclear risk of bias in five or more of the same items. Analyses were conducted for (A) anxiety/fear-related behaviour, (B) locomotor function, (C) social behaviour, and (D) cortisol levels. Data are presented as Hedges’ G standardised mean differences (SMD) and 95% confidence intervals.

## IV. DISCUSSION

Ten years after the publication of the first study of UCS conducted using zebrafish as the model animal (Piato *et al*., 2011), we performed a systematic review and meta-analysis of the literature to evaluate and synthetize the behavioural and neurochemical effects of this protocol. Despite the relatively low number of studies carried out with far fewer animals than the rodent literature, the main findings of our study show that UCS increases anxiety-like behaviour and cortisol levels while decreasing locomotor activity in zebrafish. On the other hand, no effects on social behaviour were observed in this species.

Such results somewhat correlate with the findings gathered from experiments conducted with rodents. As mentioned before, although the stress regimen is shown to consistently induce anhedonia-like behavior in rodents, several variables intrinsic to the organisms such as species, sex, age, and resilience or the protocol itself have a great impact on the outcomes measured, leading to the heterogeneity observed in the literature (Antoniuk *et al*., 2019). Results for anxiety-like behaviour (Kompagne *et al*., 2008; Cox *et al*., 2011; Zhu *et al*., 2014), locomotor function (Kumar, Kuhad & Chopra, 2011; Sequeira-Cordero *et al*., 2019), and social behaviour (Boxelaere *et al*., 2017) vary considerably depending on the conditions applied in the experiments and are still in need of a thorough systematic review to determine effect direction. The same can be said for the hormonal regulation of the stress response and related neurochemical outcomes. It is also expected to observe an increase in corticosterone and an imbalance of neurochemical markers driven by the UCS in rodents, but many reports reveal behavioural alterations in the absence of detectable modifications in these other parameters as reviewed elsewhere (Willner, 2017a; Lages *et al*., 2021).

Many factors might explain the high heterogeneity revealed between included studies and the behavioural response of fish. The number and classes of stressors used differ substantially between studies. This information is crucial since different stressors have been shown to trigger different patterns of behavioural and biochemical responses in rodents (Antoniuk *et al*., 2019). Most experiments have been conducted using mixed samples of both male and female zebrafish without reporting individualised effects of UCS by sex. Unfortunately, it is still difficult to evaluate these differential impacts since more studies are required to conduct analyses grouped by sex; however, a few experiments have already shown that stress can elicit different responses in male and female zebrafish (Rambo *et al*., 2017; Huang *et al*., 2019).

Subgroup analyses indicate that the duration of the stress protocol might also influence relevant outcomes, corroborating what was shown in previous works (Piato *et al*., 2011; Palucha-Poniewiera *et al*., 2020; Fontana *et al*., 2021). When grouping experiments by this variable, no significant effects of the stress are observed in anxiety/fear-related behaviour, locomotor function, and cortisol levels for stress regimens of up to 7 days despite the overall effects of UCS for these domains. Protocols with more than 7 days, on the other hand, show a significant effect of UCS for the same variables, indicating that regimens of more than a week of stress are necessary to reveal the deleterious consequences of stress in zebrafish. It is important to note that most experiments designed to evaluate the long-lasting effects of UCS in zebrafish were included in the group with shorter stress times. In these cases, stress sessions occur in early developmental stages and tests usually take place later in the animal’s life. This allows for a long washout period between the stress and outcome assessment that might explain the lack of effects of stress when such designs are used. Capturing UCS effects heavily depends on assessment timing (Willner, 2017a; Bosch *et al*., 2022), and tests should be scheduled to avoid observing acute effects of a single stressor as well as losing the effects of the intervention as a whole since animals are likely to eventually recover, unless the stressors coincide with a window of developmental vulnerability (Jankord *et al*., 2011).

The results of this review should be interpreted with caution considering that the main effects of the analyses were influenced by studies with a high risk of bias. Although many efforts have been made to improve the reporting quality of pre-clinical research (Sert *et al*., 2020), the publication of studies adhering to measures designed to mitigate the risk of bias associated with methodological conduct is still low (Baker *et al*., 2014; Macleod *et al*., 2015). These problems hamper the correct analysis of results and contribute to the reproducibility crisis in the biomedical field (Samsa & Samsa, 2019; Gerlai, 2019), encouraging researchers to question the validity of animal models (Worp *et al*., 2010). By excluding studies with a high risk of bias in the sensitivity analysis it was possible to visualise the direct impacts of these on distorting the main effects found in the meta-analyses for anxiety/fear-related behaviour, locomotor activity, and especially for cortisol, for which effect direction was inverted in sensitivity analysis.

In the same way, publication bias plays a part in generating misleading assumptions even in meta-analyses based on broad and rigorous systematic reviews (Worp *et al*., 2010). There is evidence of selective publishing of studies for the domains tested based on funnel plot inspection and Egger’s test evaluation, pointing to the need to conduct well-delineated experiments using this model, as these results denote a possible overestimation of the effects of chronic stress in zebrafish.

## V. CONCLUSIONS

1. The overall results of our meta-analysis reveal the effects of UCS in increasing anxiety/fear-related behaviour and cortisol levels while decreasing locomotor function;
2. No effects of stress were found for social behaviour, but the literature reporting this outcome is limited, conflicting and with evidence of bias, which warrants well-designed future experiments to fill this gap;
3. The risk of bias was considered generally high for the studies included in this review, indicating poor methodological and reporting quality of studies conducted using zebrafish;
4. We found moderate to high heterogeneity in the data, suggesting that several variables could influence the results obtained. Given the small number of studies included, it is difficult to point out the sources of variation other than the duration of the stress protocol;
5. Protocols of more than a week of stress (mostly 14 days) seem to be better suited to induce behavioural and biochemical alterations that are expected to occur with UCS;
6. The shorter duration makes zebrafish UCS protocols less time-consuming as compared with rodent protocols, which may be a convenient advantage, especially considering resource constraints;
7. Our analyses stress the need to conduct well-designed experiments using the UCS model to assess its effects on zebrafish behaviour and neurochemical parameters, further exploring the sources of variation that might influence the results, such as the nature of stressors and sex;
8. Overall, this review corroborates the need for improvement in methodological and reporting conduct across preclinical research.

## VI. AUTHOR CONTRIBUTIONS

**Matheus Gallas-Lopes:** conceptualization, data curation, formal analysis, investigation, methodology, project administration, visualisation, and writing - original draft; **Leonardo M. Bastos:** conceptualization, investigation, methodology, and writing – review & editing; **Radharani Benvenutti:** conceptualization, investigation, methodology, and writing – review & editing; **Alana C. Panzenhagen:** conceptualization, formal analysis, methodology, visualisation, and writing - review & editing; **Angelo Piato:** conceptualization, investigation, methodology, and writing – review & editing; **Ana P. Herrmann:** conceptualization, data curation, formal analysis, investigation, methodology, project administration, supervision, visualisation, and writing – review & editing;

## VIII. DATA AVAILABILITY

All data is available in Open Science Framework (https://osf.io/j2zva/).

## ACKNOWLEDGEMENTS

The authors thank the Conselho Nacional de Desenvolvimento Científico e Tecnológico (CNPq, proc. 303343/2020-6), Coordenação de Aperfeiçoamento de Pessoal de Nível Superior - Brasil (CAPES), and Pró-Reitoria de Pesquisa (PROPESQ) at Universidade Federal do Rio Grande do Sul (UFRGS) for funding and support.

