## Supplementary figures and images for "Ten years of unpredictable chronic stress in zebrafish: a systematic review and meta-analysis"

### Suppl. Fig. 1

(A) Anxiety/fear-related behaviour

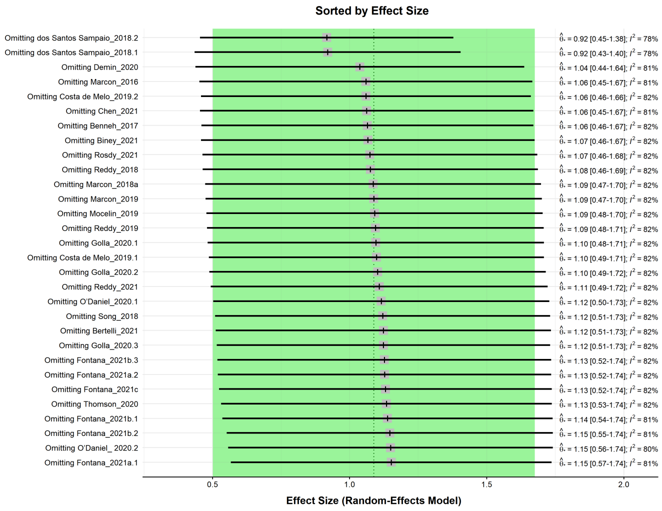


(B) Locomotor function

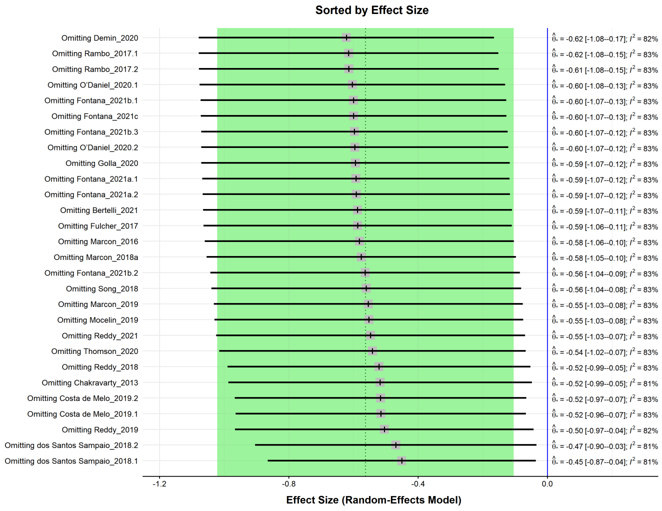


(C) Social behaviour

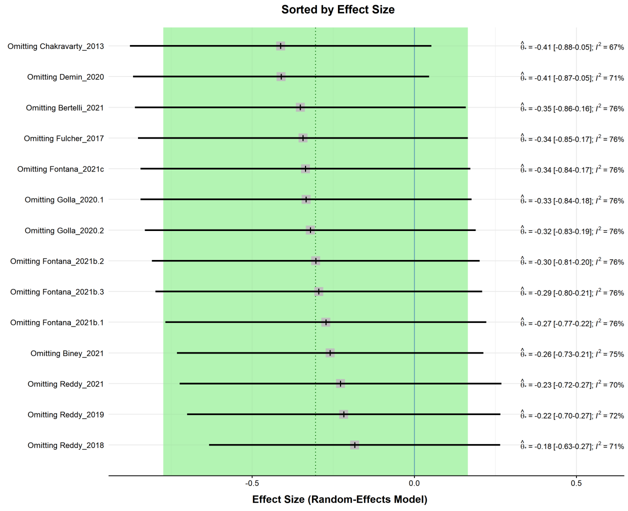


(D) Cortisol levels

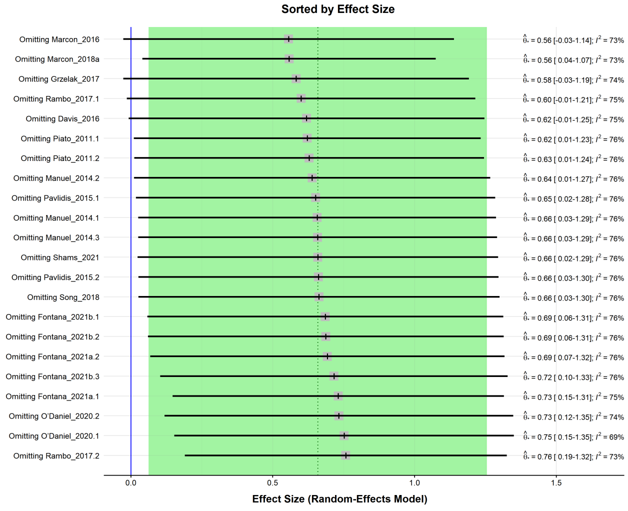
